# PINPOINT enables non-destructive nanoscopic quality control of individual lipid nanoparticles and their mRNA cargo

**DOI:** 10.64898/2026.10.11.758378

**Authors:** Pavel Banerjee, Tanmay Chatterjee, Alexander Johnson-Buck, Nicole E. Wrubel, Armin Ahnoud, Nils G. Walter

**Author notes:** These authors contributed equally: Pavel Banerjee, Tanmay Chatterjee.

## Abstract

Quality control of lipid nanoparticles (LNPs) is central to development of mRNA therapeutics, yet ensemble measurements obscure critical heterogeneity in particle morphology and mRNA cargo loading. Here we introduce PINPOINT (Profiling Individual Nanoparticles with Precise Observation of INternalized Therapeutics), a single-particle nanoscopy platform for non-destructive, multiparametric profiling of preformulated LNP samples. Super-resolution imaging with Nile Red resolves size and morphology, while an optimized permeabilization strategy provides selective access to encapsulated mRNA without compromising LNP integrity. Single-particle fluorescence measurements, including RiboGreen and sequence-specific probes, enable quantification and validation of mRNA loading and identity. Comparative analyses of bare and encapsulated mRNA reveal kinetic and structural features, exposing heterogeneity masked by bulk assays. By integrating structural and molecular readouts at the single-particle level, PINPOINT establishes a quantitative framework for assessing of LNP composition and payload distribution, offering actionable insights for improving batch consistency and guiding the rational design of next-generation mRNA delivery systems.

---

Messenger RNA (mRNA) therapeutics have transformed modern medicine, enabling vaccines, protein replacement and gene therapy^1–3^. Their clinical success depends on efficient mRNA encapsulation and delivery, a challenge addressed by lipid nanoparticles (LNPs) that underpin FDA-approved nucleic acid medicines, including siRNA drugs and several COVID-19 vaccines^4,5^. Stable, potent LNPs are typically formulated from ionizable lipids, phospholipids, cholesterol and polyethylene glycol conjugates^6–8^. The performance of these LNP-based therapeutics is highly sensitive to formulation and manufacturing variables, including microfluidic mixing and pH-driven assembly^9–14^. While advances in production have improved control over size and morphology, they also underscore how strongly efficacy, toxicity, and biodistribution depend on physicochemical attributes, making rigorous characterization essential for formulation development^15–20^.

Routine LNP analytics remain dominated by bulk measurements such as dynamic light scattering (DLS), nanoparticle tracking analysis (NTA) and the Quant-iT RiboGreen assay, which report population averages and cannot resolve particle-to-particle variability in size, shape or cargo encapsulation^21,22^. Thus, these methods can mask heterogeneities, such as the presence of substantial fractions of empty, sparsely loaded particles, that may influence therapeutic outcomes^22–24^. Electron microscopy provides high-resolution structural information but typically requires sample preparation that perturbs native LNP states and complicates structure-function analysis, and cannot report on the sequence of any RNA cargo^19,25^. Fluorescence-based single-particle approaches can distinguish loaded from empty particles^23,27^, but stable labelling may impact loading or efficacy and thus may be incompatible with release testing and scale up of the particles, and intensity-based readouts rarely report on cargo integrity or precise sequence identity^26–28^. Critically, direct access to encapsulated mRNA often requires disruptive processing that compromises LNP structure and obscures relationships between morphology, loading and integrity^20,23,27^. These limitations motivate non-destructive platforms that can jointly quantify morphology and cargo loading, identity and integrity in individual LNPs.

To address these gaps, we developed PINPOINT (Profiling Individual Nanoparticles with Precise Observation of INternalized Therapeutics), a single-molecule nanoscopy platform for multiparameter profiling of individual LNPs, including preformulated samples with defined mRNA cargo compositions. PINPOINT combines super-resolution Nile Red imaging for sizing and morphology with controlled Triton X-100 (TX-100) permeabilization that enables probe access to encapsulated mRNA while preserving measurable particle morphology. Single-particle RiboGreen imaging and sequence-specific fluorescent probes quantify mRNA loading and assess cargo identity and regional accessibility within surface-captured particles. By linking particle morphology with single molecule kinetic profiling from bare and encapsulated mRNA, PINPOINT reveals heterogeneity masked by bulk methods, enabling quantitative quality control to support batch comparison, formulation optimization and the rational design of next-generation mRNA delivery systems.

## Fingerprinting lipid nanoparticles for quality control

To meet the need for a non-destructive, amplification-free approach that reads out multiple orthogonal attributes from preformulated LNP samples without covalent modification of their components, we developed PINPOINT, a nanoscopic fingerprinting platform implemented on a commercial total internal reflection fluorescence (TIRF) microscope^29–32^. The workflow addresses quality attributes relevant to mRNA–LNP formulation and manufacturing, progressing from particle morphology to cargo quantification and identity (Fig. 1a). PINPOINT immobilizes LNPs on a passivated glass coverslip via anti-PEG capture without covalent modification^33^. Surface-captured LNPs are then interrogated using three complementary modalities (Fig. 1a, b): super-resolution Nile Red imaging to assess LNP size and morphology^34,35^, single-particle RiboGreen fluorescence to quantify RNA content as a proxy for mRNA copy number^36^, and sequence-specific fluorescent oligonucleotide probes for highly specific identification of cargo through repeated transient binding that yields a kinetic fingerprint distinct from nonspecific interactions^31,32^. Probe and capture sequences are listed in Supplementary Table 1.

**Fig. 1.**
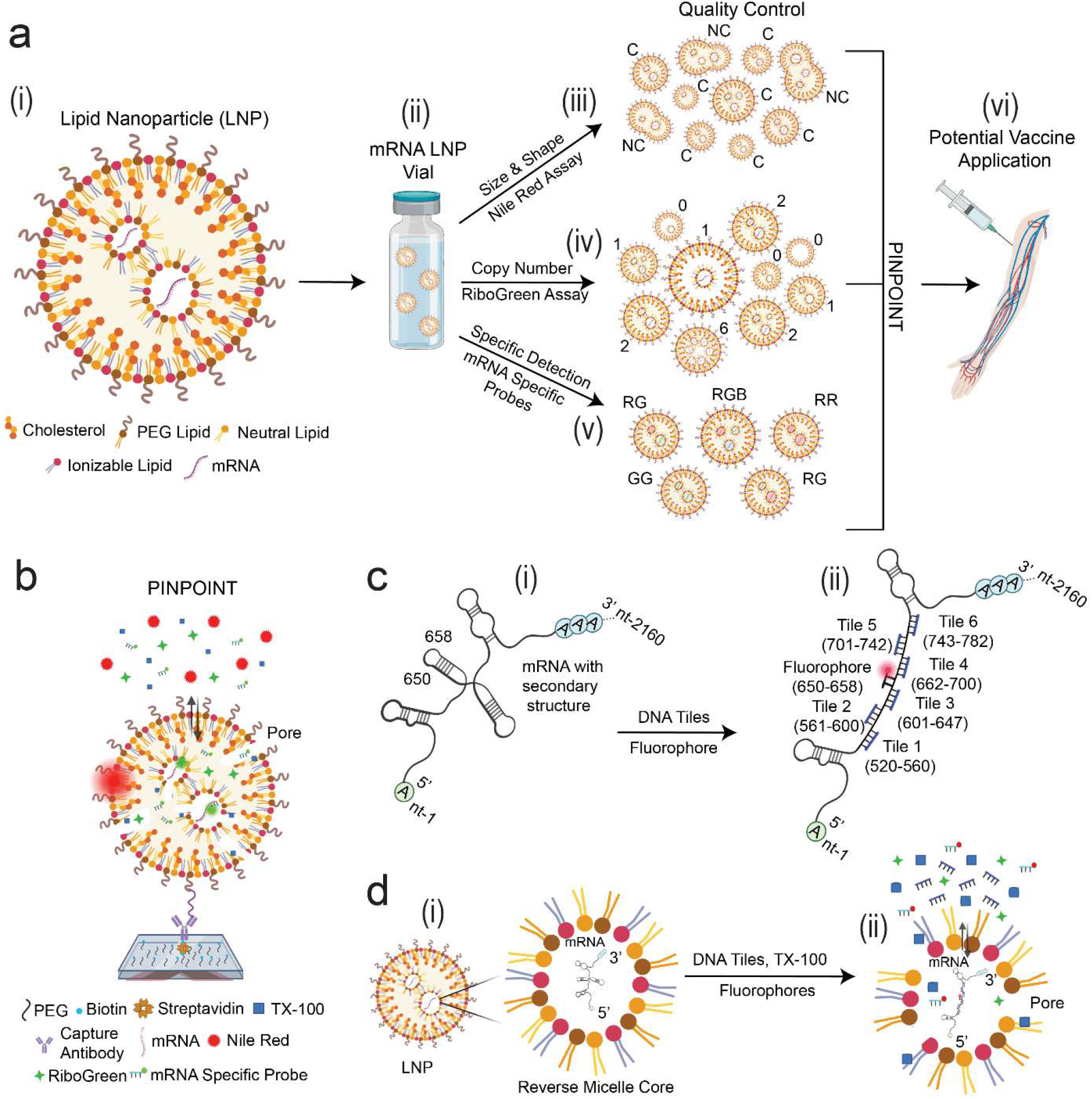
PINPOINT fingerprinting workflow enables single-platform quality control for mRNA lipid nanoparticles. **a,** Overall workflow for lipid nanoparticle (LNP) quality control using PINPOINT. (i) Schematic of an LNP and its basic constituents: cholesterol, PEG lipid, neutral lipid, ionizable lipid, and mRNA (cargo). (ii) Illustration of an mRNA LNP therapeutic vial. (iii–vi) Quality control checks performed in a single platform: (iii) Determination of size and shape of individual LNPs via the Nile Red assay; (iv) Estimation of mRNA (cargo) copy numbers per LNP by RiboGreen assay; (v) Specific mRNA (cargo) detection inside LNPs using mRNA-specific probes; (vi) Potential LNP therapeutic administration following all quality control checks. **b**, Nanoscopic characterization of surface-captured LNPs with Nile Red, RiboGreen, and/or mRNA-specific probes. TIRF microscopy measures both intensity signatures from static probe interactions (RiboGreen) and dynamic signatures from repeated probe binding and dissociation (Nile Red and mRNA-specific kinetic fingerprinting probes) for nanoscopic morphology determination and highly specific sequence detection. **c**, (i) Schematic representation of studied mRNAs (approximately 2100 nt) target region for probing at nt 650–658 that may be occluded by secondary structure. (ii) This accessibility issue is mitigated by a DNA tiling strategy (three ∼40 nt tiles flanking each side of the target site, or six tiles in total) that increases accessibility of the target site by disrupting secondary structure and enhances detection by mRNA-specific probes. **d**, Detection of encapsulated mRNA inside the reverse micelle core of LNPs (i) with secondary structure prior to detection, and (ii) after DNA tiling and permeabilization using TX-100, allowing specific detection of encapsulated mRNA.

We developed and validated PINPOINT using three externally prepared LNP formulations (LNP1001, LNP1002, and LNP1004) with distinct mRNA cargo compositions containing mRNA1, mRNA2, or both cargos (Supplementary Table 2), alongside a commercially sourced cargo-negative LNP formulation (LNP-ALC0315) as an RNA-negative control (Supplementary Fig. 1). Cargo identification is challenging for conventional *in situ* hybridization approaches because the mRNAs share ∼98% sequence identity. To increase sequence discrimination, we used a kinetic fingerprinting readout based on transient binding of short probes (8–9 nt), whose interaction kinetics are highly sensitive to small differences in sequence^31^. To minimize perturbation of LNPs, we further opted for isothermal *in situ* hybridization under mild conditions; however, a significant obstacle arises from the substantial secondary structure across the ∼2.1-kb transcripts, which can occlude binding sites and introduce structure-dependent heterogeneity^37^. We therefore targeted a short window (nt 650–658) that would allow discrimination between the two closely related mRNA cargos and used a local DNA-tiling strategy, placing three unlabelled, fully complementary 40-nt tiles upstream and three downstream of the probed site to reduce local folding and stabilize probe access (Fig. 1c and Supplementary Fig. 2)^38^. Because cargo is sequestered within the LNP interior, we paired tiling with a mild, optimized TX-100 permeabilization that permits probe entry while preserving overall particle contents and morphology (Fig. 1d)^39,40^. This assay design forms the foundation for the remaining sections, in which we quantify LNP morphology, map a non-destructive permeability window, and identify and interrogate encapsulated mRNA at the single-particle level.

## Super-resolution imaging reveals LNP size and morphology

Particle size is a critical quality attribute for LNP performance and is typically reported as a hydrodynamic diameter by DLS^29,30^, but as an ensemble technique DLS has limited sensitivity to minor subpopulations and does not directly report on particle morphology^19^. We therefore implemented a super-resolution module within PINPOINT to quantify LNP size and morphology directly from individual particles using Nile Red, a fluorogenic dye whose emission is enhanced in hydrophobic environments, based on the PAINT technique^34,35^. Repeated association and dissociation of Nile Red at the LNP interface produces temporally sparse fluorescence bursts that enabled particle detection and nanoscopic reconstruction from localizations of individual binding events (Fig. 2a)^31,32^. LNPs were surface-captured via an anti-PEG antibody on passivated coverslips^33^ and analyzed using localization-based reconstruction (Methods; Supplementary Figs. 3, 4)^31,32,34^. Assay conditions were optimized to preserve sparse-event kinetics and minimize background (Extended Data Fig. 1a, b). Nile Red imaging yielded bright puncta for captured LNPs, whereas no-LNP blank controls, containing Nile Red and the same surface chemistry but no added LNPs, exhibited only sparse short-lived events with weak signal, likely due to residual surface binding of the dye (Fig. 2b and Extended Data Fig. 1c). For each bright particle accepted, localizations were compiled into a super-resolution reconstruction (Fig. 2c), from which we extracted particle-resolved diameter, area and circularity index^34^. Particle diameter distributions from LNP samples showed far more particles than blank measurements (Fig. 2d), suggesting that the majority of signal arises from LNPs rather than background binding or Nile Red aggregates. Within single fields of view, PINPOINT resolved distinct size classes consistent with subpopulations masked in bulk measurements (Extended Data Fig. 1d).

**Fig. 2.**
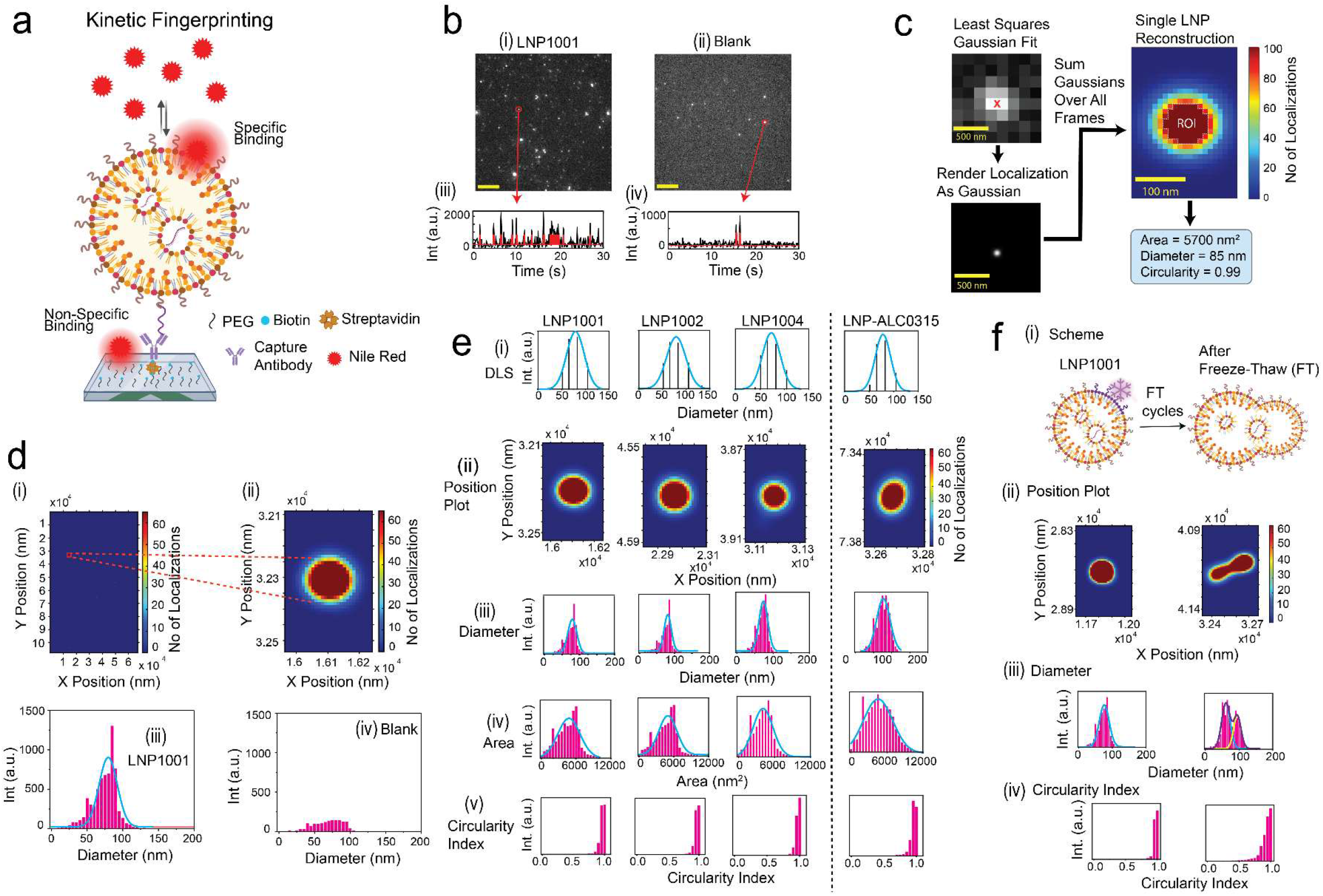
Fluorescence nanoscopy enables precise analysis of LNP size, shape, and morphology. **a,** PAINT nanoscopy of surface-captured LNPs with TIRF observation of the reversible binding of Nile Red enables size and morphology measurements. **b**, Average movie frames of a representative TIRF microscope field of view (FOV; scale bar, 5 µm) showing bright puncta where Nile Red probes bind at or near the imaging surface: representative FOVs shown for (i) LNP1001 and (ii) a no-LNP blank control. Representative kinetic fingerprinting traces from (iii) LNP1001 and (iv) a no-LNP blank control are also shown and comprise raw intensity-versus-time traces (black lines) idealized by hidden Markov modeling (red lines). Temperature: 25 °C; Acquisition time: 5 min/FOV; Probe: Nile Red at 1 nM. **c**, Workflow for super-resolution analysis: spots from Nile Red binding to individual LNP particles are localized using least squares Gaussian fitting; localizations are represented as Gaussian intensity distributions and accumulated across all movie frames to enable reconstruction of single LNPs and precise determination of size and shape. **d**, Localization reconstructions showing (i) an entire FOV of LNPs and (ii) a zoomed-in view of a single LNP; (iii) histogram of size estimates for LNP1001 (background-subtracted) across one FOV with Gaussian fit; (iv) corresponding histogram for a no-LNP blank control. **e**, Profiling LNP1001, LNP1002, LNP1004, and LNP-ALC0315: (i) dynamic light scattering (DLS) diameter distribution; (ii) reconstructions of individual LNPs; estimation of (iii) diameter, (iv) area, and (v) shape (circularity index) of distinct LNPs from super-resolved images across the FOV. **f**, Assessment of freeze–thaw (FT) effects on LNP1001: (i) schematic illustration of structural changes; (ii) reconstructions; (iii) diameter histograms; (iv) circularity index distributions after FT treatment.

We applied this assay to four LNP batches (LNP1001, LNP1002, LNP1004, and LNP-ALC0315). PINPOINT mean diameters agreed closely with DLS while additionally providing particle-level morphology distributions and representative super-resolution reconstructions of individual particles (Fig. 2e and Supplementary Table 3; Extended Data Fig. 1e), which were predominantly circular under these conditions. To test sensitivity to handling-induced structural changes^30,41^, we subjected LNP1001 to repeated freeze–thaw cycling between liquid nitrogen and a 37°C water bath. Although untreated LNP1001 particles already showed some shape heterogeneity, with circularity indices distributed below the ideal value of 1.0, freeze–thaw stress broadened this distribution and increased the fraction of visibly non-circular particles in super-resolution reconstructions. DLS and PINPOINT both reported a shift from an apparently unimodal to a bimodal size distribution (Fig. 2f, Extended Data Fig. 1f), while PINPOINT additionally provided particle-resolved shape information, revealing an emergent non-circular subpopulation that was not apparent from DLS alone (Fig. 2f and Supplementary Table 3).

## Quantifying LNP permeability and cargo access

Bulk assays can estimate RNA content in LNP formulations, but they typically rely on detergent-mediated lysis, precluding direct measurement of encapsulated mRNA accessibility while retaining particle-level morphological information^36,39^. To enable non-destructive cargo access at the single-LNP level, we incorporated a controlled permeabilization module into PINPOINT using mild surfactant (TX-100) conditions, together with fluorescence readouts of RNA accessibility (Fig. 3a). By operating below the critical micelle concentration (CMC) of TX-100—i.e., at concentrations of 0.005–0.01% v/v—we aimed to modulate permeability without driving full micellar solubilization^39,40^, and used RiboGreen as a static-access reporter complementary to a kinetic readout from mRNA-specific probes. We first defined a TX-100 operating window that increased RNA accessibility while preserving LNP morphology. In bulk spectra of LNP1001 with RiboGreen, fluorescence rose sharply over a narrow TX-100 range, consistent with a transition from an essentially impermeable state to one in which encapsulated RNA is dye-accessible (Fig. 3b and Extended Data Fig. 2a). This transition guided subsequent experiments within the 0.005– 0.01% regime, centered at 0.008% v/v. Under matched imaging conditions, RiboGreen TIRF revealed puncta only within this window, indicating surfactant-assisted access to internal RNA (Fig. 3c (ii, iv)). Importantly, Nile Red super-resolution sizing showed no significant change in LNP diameter across optimized TX-100 concentrations, whereas higher surfactant levels produced disruption consistent with dissolution, leading us to avoid these levels (Fig. 3c). Thus, TX-100 can be tuned to increase cargo accessibility while maintaining particle morphology at the resolution of our nanoscopy assay.

**Fig. 3.**
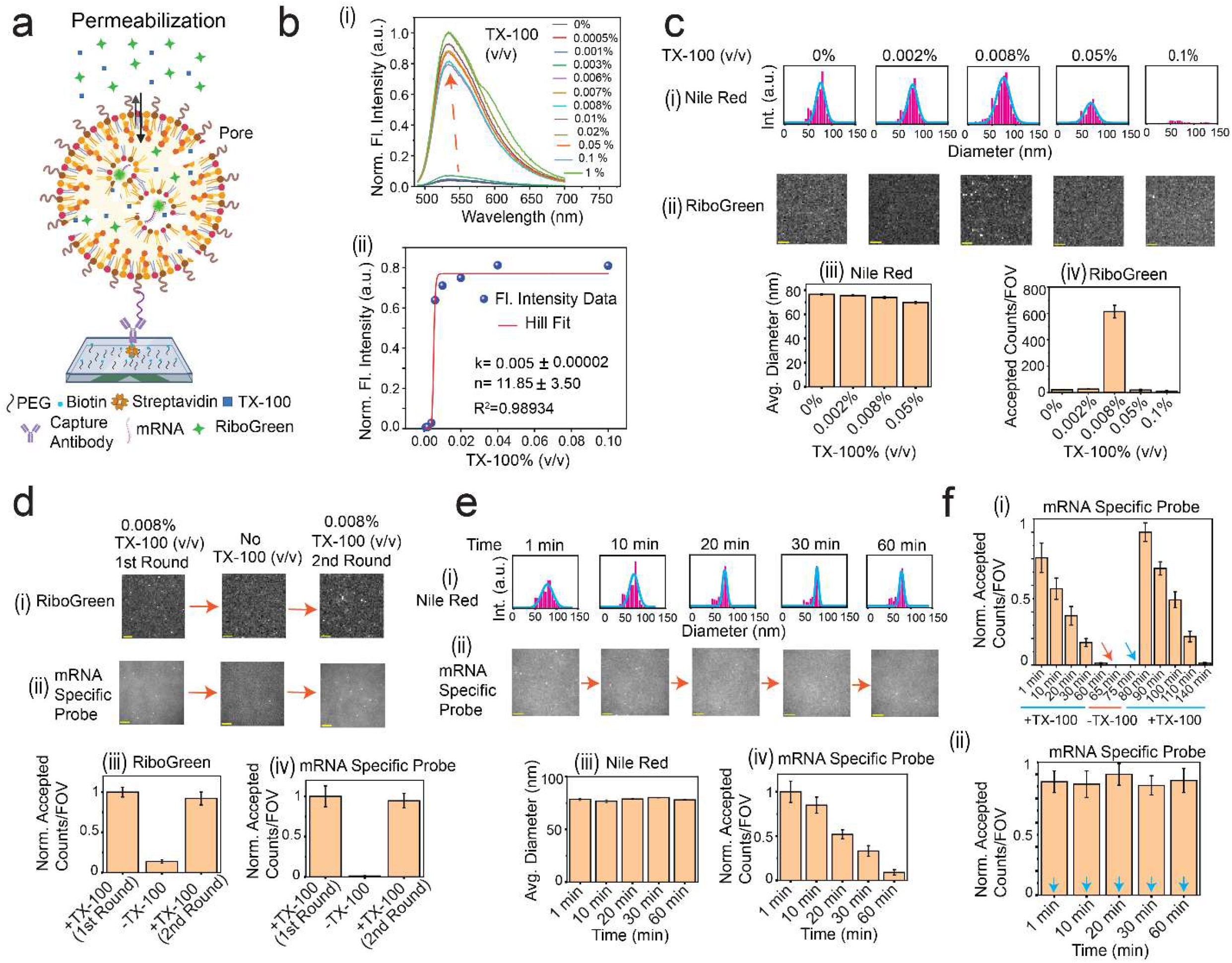
Quantitative assays reveal LNP permeability and cargo access via controlled membrane permeabilization. **a,** Nanoscopic fingerprinting of surface-captured LNPs with RiboGreen enables cargo access through static probe binding, after permeabilization using optimized TX-100 concentration. Fingerprints are measured by TIRF microscopy. **b**, (i) Steady-state fluorescence spectra (normalized fluorescence intensity vs wavelength) of LNP1001 with RiboGreen at 0.5% across a range of TX-100 concentrations (0%–1% v/v), highlighting the membrane permeabilization sensitivity window. (ii) Hill fit of steady-state fluorescence intensity data characterizing the TX-100 concentration response. **c**, (i) Size profiling of LNP1001 by Nile Red assay, showing diameter distributions from reconstructed images with varying TX-100 concentrations. (ii) RiboGreen imaging frames for mRNA1 accessibility inside LNP1001 under different TX-100 concentrations (scale bar, 5 µm). (iii) Changes in average LNP diameter with TX-100 concentration (Nile Red assay). (iv) Accepted counts per field of view (FOV) for RiboGreen assay quantification with varying TX-100 concentrations. Temperature: 25 °C; Acquisition time: 15 s/FOV; Probe: Nile Red at 1 nM; RiboGreen at 0.5%. **d**, Cargo accessibility of mRNA in LNPs in the presence (first and second rounds) and absence of 0.008% TX-100. (i) RiboGreen assay imaging; (ii) mRNA-specific probe binding assay; (iii) Quantified, normalized accepted counts per FOV via RiboGreen assay; (iv) Quantified counts per FOV via mRNA-specific probe binding. **e**, Changes in LNP1001 size and cargo accessibility in the presence of 0.008% TX-100: (i) Nile Red assay diameter distributions over time; (ii) TIRF imaging frames of mRNA1-specific probe binding (scale bar, 5 µm); (iii) Average diameters over time using Nile Red data; (iv) Quantification of accepted counts per FOV for mRNA1-specific probes over time. Temperature: 25 °C; Acquisition time: 5 min/FOV; Probe: Nile Red at 1 nM; mRNA1 FP (650–658) _Middle Region_9 nt_Cy5 at 50 nM. **f**, (i) Quantification of mRNA1-specific probe accessibility inside LNP1001 over time (0–60 min) with TX-100 removal at 60 min (red arrow), followed by addition of fresh 0.008% TX-100 buffer (cyan arrow). (ii) Quantification of probe accessibility over time with successive external additions of fresh 0.008% TX-100 buffer at each time point (cyan arrows). Acquisition time: 5 min/FOV; Probe: mRNA1 FP (650–658) _Middle Region_9 nt_Cy5 at 50 nM; DNA tiles at 6 nM. Black error bars indicate standard errors of the mean from three independent replicates.

Cargo access was reversible and could be toggled by TX-100 availability: removing TX-100 abolished RiboGreen signals, as well as those of mRNA specific probes, and re-addition restored accessibility (Fig. 3d). Under static conditions, accessibility decayed over time across the operational window, as RiboGreen and mRNA-probe signals decreased while Nile Red-detected particle counts and size distributions remained stable (Fig. 3e and Extended Data Fig. 2b–d). DNA tile addition did not recover accessibility (Extended Data Fig. 2e), whereas replenishing surfactant by buffer exchange followed by fresh TX-100 or by repeated TX-100 addition restored and sustained access (Fig. 3f). These results support a sub-CMC regime in which TX-100 transiently increases permeability, but effective access declines over time due to surface adsorption^39,40^.

## Sequence specific detection of mRNA cargo

Sequence-specific single-molecule detection of intact mRNA is challenged by transcript length and secondary structure, which can occlude short probe binding sites and create time-dependent accessibility^30^. To demonstrate solutions for this challenge, we first established sequence-targeted detection on bare mRNA before moving to encapsulated cargo. We immobilized mRNA1 on passivated coverslips and interrogated a defined window (nt 650–658, Supplementary Fig. 2) with a short, Cy5-labelled 9-nt probe (Fig. 4a). Binding produced intensity time traces with repeated association and dissociation, which were idealized by hidden Markov modelling and filtered using kinetic criteria to reject nonspecific interactions (Methods; Fig. 4b and Extended Data Fig. 3a, general parameter sets in Supplementary Table 4)^31,32,42,43^. Under these conditions, blank chambers yielded near-zero (2.33 ± 0.47) accepted traces, supporting specific detection. However, without tiles, probe binding to mRNA1 was strongly intermittent, with burst-like periods of frequent events separated by extended intervals with little or no binding (Fig. 4c and Extended Data Fig. 3b), consistent with structure-gated switching between probe-accessible and probe-inaccessible states^43^. Fitting the cumulative distribution of burst durations yielded an accessibility transition rate constant (k_acc→inacc_ = 0.027 s^-1^), suggesting that the underlying structural dynamics occur on a timescale of tens of seconds (Extended Data Fig. 3c).

**Fig. 4.**
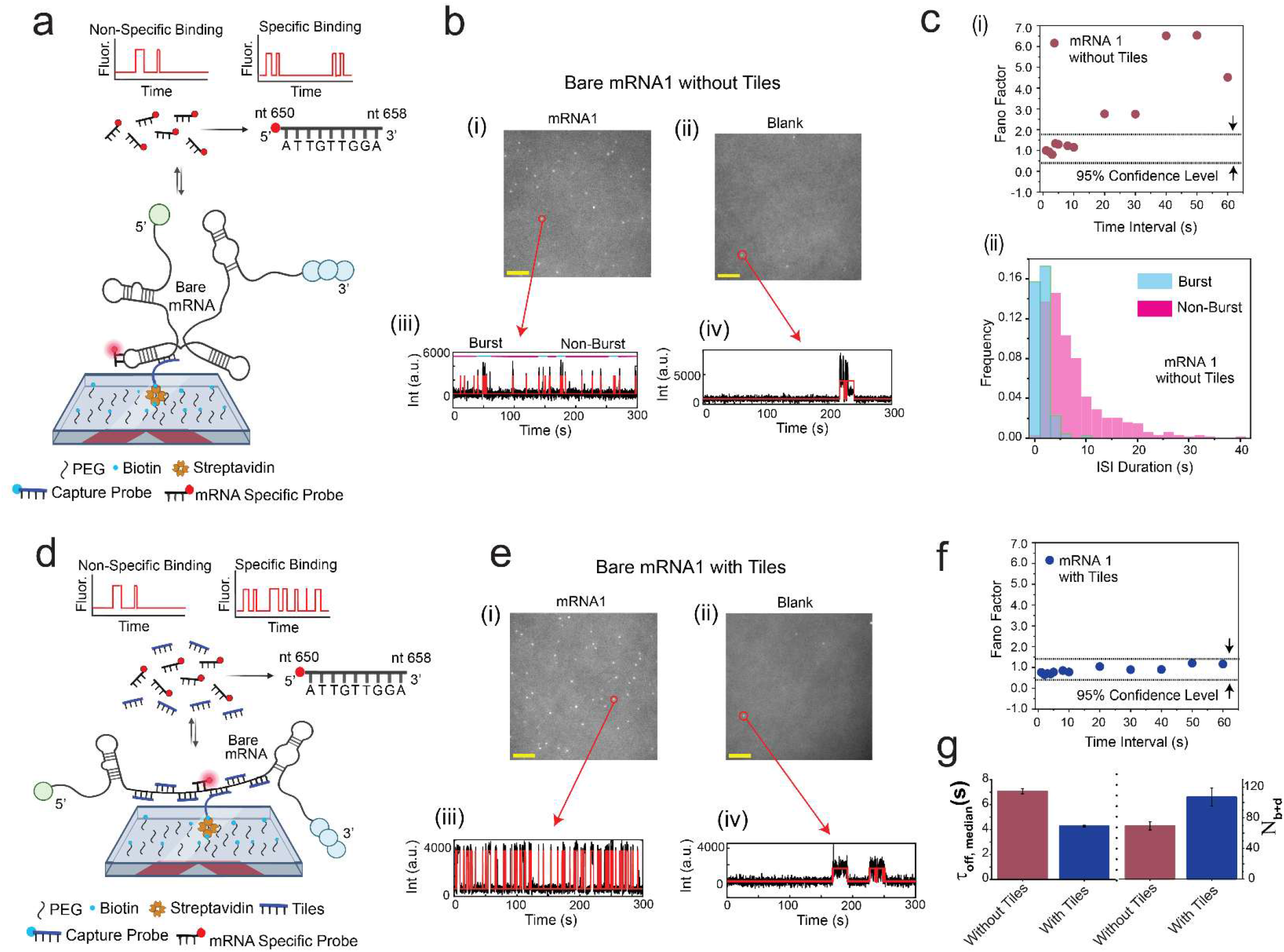
Sequence-targeted single-molecule detection of bare mRNA1 with and without DNA tiling. **a,** Nanoscopic fingerprinting of surface-captured bare mRNA1 (without tiles) using mRNA-specific probes reveals diverse temporal patterns of repeated binding and dissociation, distinct from nonspecific probe interactions. Fingerprints are measured by TIRF microscopy. **b**, Average movie frames of a representative TIRF microscope field of view (FOV; scale bar, 5 µm) showing bright puncta where specific probes bind at or near the imaging surface: (i) representative bare mRNA1 and (ii) blank sample. Representative kinetic fingerprinting traces of (iii) bare mRNA1 and (iv) blank sample—raw intensity-versus-time traces (black lines) idealized by hidden Markov modeling (red lines) to extract kinetic parameters. All data are without tiles. **c**, (i) Fano factor analysis calculated across various time intervals for specific probe–bare mRNA1 interaction (without tiles), indicating non-Poissonian statistics (burgundy circles); dashed lines show the 95% confidence interval for Poisson behaviour. (ii) Cumulative histogram of interspike interval (ISI) duration during burst (cyan) and non-burst (magenta) periods for mRNA1 without tiles. **d**, Nanoscopic fingerprinting of surface-captured bare mRNA1 (with tiles) using mRNA-specific probes reveals diverse temporal patterns of repeated binding and dissociation. Fingerprints are measured by TIRF microscopy. **e**, Average movie frames of a representative TIRF microscope FOV (scale bar, 5 µm) showing probe binding at or near the surface: (i) bare mRNA1 and (ii) blank sample. Representative kinetic fingerprinting traces: (iii) bare mRNA1 and (iv) blank sample; traces idealized by hidden Markov modeling. All data are with tiles. **f**, Fano factor analysis across time intervals for probe–bare mRNA1 interaction with tiles, showing Poissonian statistics (blue circles), with dashed lines indicating the 95% confidence interval for Poisson behaviour. **g**, Comparison of τ_off,median_ and N_b+d_ for bare mRNA1–probe interactions through nanoscopic fingerprinting without (burgundy) and with (blue) tiles. Temperature: 25 °C; Acquisition time: 5 min/FOV; Probe: mRNA1 FP (650–658) _Middle Region_9 nt_Cy5 at 50 nM; DNA tiles at 6 nM. Black error bars indicate standard errors of the mean from three independent replicates.

To increase accessibility of the probe binding site, we implemented a local DNA-tiling strategy in which six unlabeled ∼40-nt DNA oligonucleotides (tiles) adjacent, both upstream and downstream, to the probed site to disrupt competing secondary structure (Fig. 4d, Supplementary Fig. 2). As expected for a suppressed local mRNA secondary structure, tiling increased the frequency and temporal uniformity of specific probe binding and increased the number of accepted trajectory counts per FOV ∼5-fold while maintaining low background (Fig. 4e and Extended Data Fig. 3d). Tile assembly affected assay performance, with thermal cycling and longer incubation increasing accepted counts, and pre-capture outperforming co-capture (Extended Data Fig. 3e; Supplementary Fig. 5). Tiling also suppressed the bursting behavior, yielding a Fano factor consistent with Poisson binding statistics and reducing unbound times (Fig. 4f, g and Extended Data Fig. 3f)^43,44^. Together, these results establish that short probes combined with kinetic fingerprinting can achieve sequence-targeted single-molecule detection of structured mRNAs, and that local tiling eliminates structure-gated binding to yield a more uniform, quantitative assay suitable for downstream cargo identification in LNPs.

## Probing encapsulated mRNA identity and regional accessibility

A key need in mRNA–LNP analytics is to assess encapsulated cargo identity and integrity *in situ* without destroying the particle and, with it, any information about population heterogeneity. Using the non-destructive permeability window defined above, we combined sub-CMC TX-100 (0.008% v/v) with sequence-specific kinetic fingerprinting probes to detect encapsulated mRNA in morphology-preserved, surface-captured LNPs by TIRF microscopy (Fig. 5a). Specificity derives from repeated binding and dissociation patterns with kinetic filtering to reject nonspecific interactions (Methods)^32,42^. We first targeted encapsulated mRNA1 in LNP1001 at the internal binding site (nt 650–658). Without tiles, probe binding was detectable but strongly intermittent, with burst-like periods separated by long inactive intervals (Fig. 5b (i, iii) and Extended Data Fig. 4a, b), consistent with structure-gated switching between probe-accessible and probe-inaccessible states. Encapsulation accentuated this gating relative to the bare mRNA1 (Extended Data Fig. 4d - Introducing local DNA tiling regularized access: tiled, encapsulated mRNA1 produced denser event trains (Fig. 5b (ii, iv)), binding statistics approached Poisson behaviour (Fig. 5b (v)), and kinetic fingerprints shifted toward more frequent interrogation (Fig. 5b (vi) and Extended Data Fig. 4c)^43^. Comparing tiled mRNA1 in bare versus encapsulated contexts further showed that encapsulation altered both bound and unbound time distributions, yielding shorter τ_on,med_ and longer τ_off,med_ than bare mRNA1 (Fig. 5c), consistent with both reduced binding site accessibility and destabilization of hybridization inside LNPs containing both zwitterionic and ionizable cationic lipids.

**Fig. 5.**
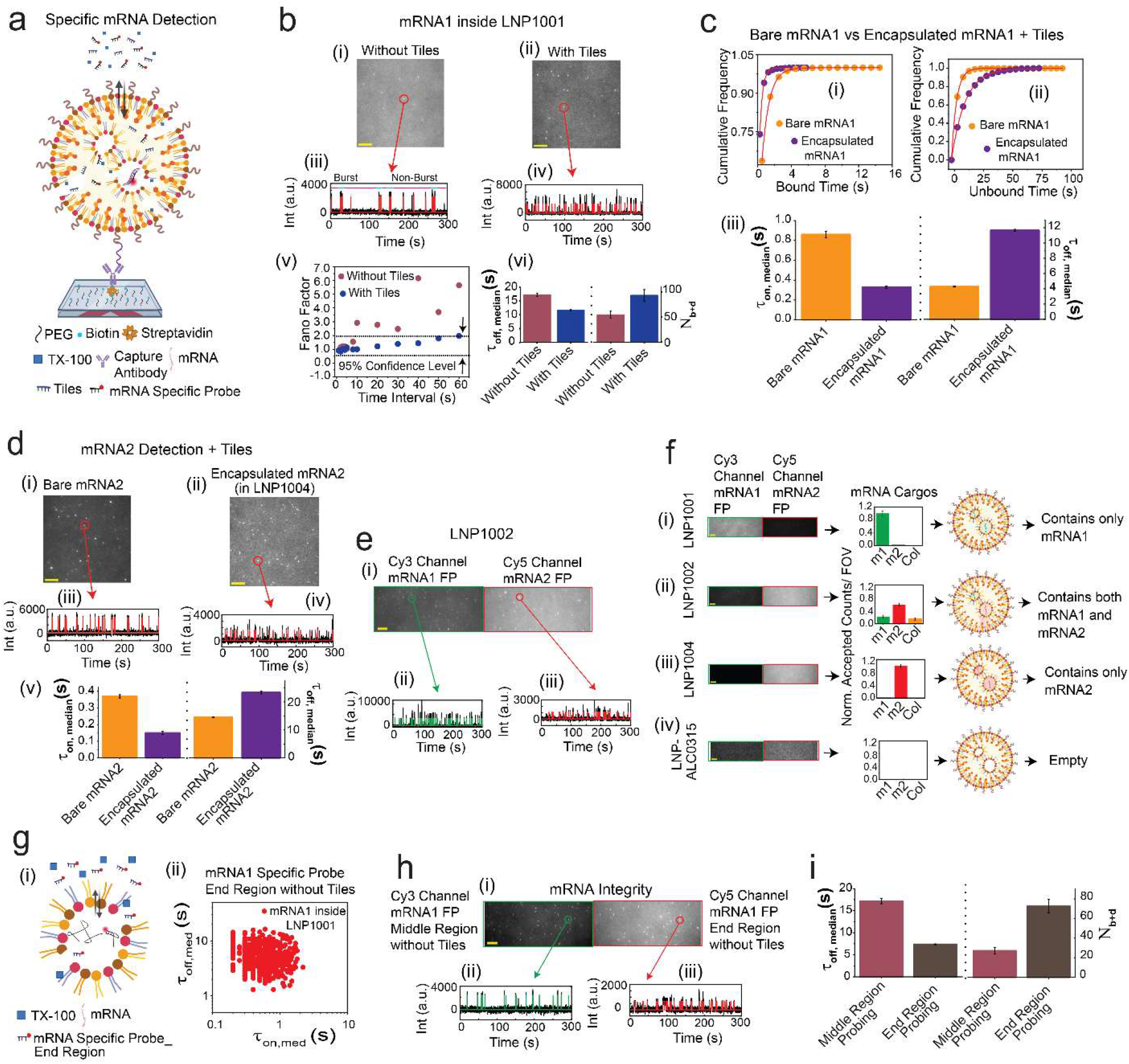
Specific single-molecule assays enable sequence-specific and multi-region probing of encapsulated mRNA within LNPs. **a,** Nanoscopic fingerprinting of surface-captured LNPs with mRNA-specific probes enables access to encapsulated cargos through diverse temporal binding and dissociation patterns, following permeabilization with optimized TX-100 concentration. Detection is performed by TIRF microscopy. **b**, Average movie frames of a representative TIRF microscope field of view (FOV; scale bar, 5 µm) showing probe binding to encapsulated mRNA1 inside LNP1001: (i) without tiles and (ii) with tiles. Representative kinetic fingerprinting traces of (iii) without tiles and (iv) with tiles—intensity-versus-time traces (black lines) idealized by hidden Markov modeling (red lines). (v) Fano factor analysis across time intervals for probe–encapsulated mRNA1 interactions: non-Poissonian statistics (burgundy circles, without tiles), Poissonian statistics (blue circles, with tiles), with dashed lines indicating the 95% confidence interval for Poisson behaviour. (vi) Comparison of τ_off,median_ and N_b+d_ for encapsulated mRNA1–probe interactions without (burgundy) and with (blue) tiles. **c**, Cumulative dwell time analyses for bare and encapsulated mRNA1 with tiles: (i) bound and (ii) unbound time histograms with τ_on,med_ and τ_off,med_ determined by single-exponential fitting; (iii) comparison of τ_on,median_ and τ_off,median_ for bare (yellow) versus encapsulated mRNA1 (purple). Temperature: 25 °C; Acquisition time: 5 min/FOV; Probe: mRNA1 FP (650–658) _Middle Region_9 nt_Cy5 at 50 nM; DNA tiles at 6 nM; TX-100: 0.008%. **d**, Detection of bare and encapsulated mRNA2 with tiles: (i) and (ii) movie frames of bare mRNA2 and encapsulated mRNA2 (in LNP1004); (iii) and (iv) kinetic fingerprinting traces; (v) comparison of τ_on,median_ and τ_off,median_ for bare (yellow) and encapsulated mRNA2 (purple). Temperature: 25 °C; Acquisition time: 5 min/FOV; Probe: mRNA2 FP (650– 660) _Middle Region_11 nt_Cy5 at 50 nM; TX-100: 0.008%. **e**, Dual-channel detection of mRNA encapsulated inside LNP1002: (i) TIRF movie FOV divided into Cy3 (mRNA1) and Cy5 (mRNA2) channels; (ii) and (iii) representative kinetic fingerprinting traces of encapsulated mRNA1 and mRNA2. **f**, Two color experiment for multiplexed cargo detection in LNP1001, LNP1002, LNP1004, and LNP-ALC0315 (i-iv): Cy3 channel (mRNA1) and Cy5 channel (mRNA2) detection show accepted counts per FOV and schematic of corresponding LNP cargo profiles. **g**, (i) Schematic for end-region probing of encapsulated mRNA by specific probe. (ii) Scatterplots of dwell time analysis (τ_on,med_ vs τ_off,med_) for encapsulated mRNA1 in LNP1001 with end-region probe. **h**, Two-channel experiments for mRNA1 integrity analysis inside LNP1001: (i) Cy3 channel probing middle region; Cy5 channel probing end region. Representative kinetic fingerprinting traces for (ii) middle region and (iii) end region (raw traces idealized with hidden Markov modeling). All experiments without DNA tiles. **i**, Comparison of τ_off,median_ and N_b+d_ for mRNA1 integrity probing through middle region (burgundy) and end region (brown). Temperature: 25 °C; Acquisition time: 5 min/FOV; Target: LNP1001 at 100 pM; Probe: mRNA1 FP (650–658) _Middle Region_9 nt_Cy3 at 50 nM; mRNA1 FP (2033–2040) _End Region_9 nt_Cy5 at 50 nM; DNA tiles at 6 nM; TX-100: 0.008%. Black error bars indicate standard errors of the mean from three independent replicates.

This behavior generalized to a second sequence. Using an 11-nt probe against an internal binding site (nt 650–660) of mRNA2, PINPOINT detected both bare and LNP-encapsulated mRNA2 and revealed similar dependence of kinetics on LNP encapsulation as with mRNA1 (Fig. 5d and Supplementary Fig. 6). We then leveraged the short-probe strategy for multiplexed cargo identification. Dual-color probing enabled simultaneous detection of mRNA1 and mRNA2 within the same field of view (Fig. 5e). In a multiplexed detection experiment using three LNP formulations and an independent cargo-negative LNP-ALC0315 control, two-channel sequence-specific kinetic fingerprinting detected the expected cargo compositions: LNP1001 produced only mRNA1-specific signals, LNP1002 produced both mRNA1- and mRNA2-specific signals, LNP1004 produced only mRNA2-specific signals, and the LNP-ALC0315 control produced no accepted mRNA-specific signal above background (Fig. 5f). Colocalization analysis of LNP1002 suggests co-detection of both mRNAs in only a minority of particles (∼25%), consistent with an approximate independent Poisson-loading model with an average of ∼1 mRNA per LNP and with approximately twice as many accepted mRNA2-specific signals as mRNA1-specific signals under the assay conditions (Fig. 5f).

Toward developing a multi-region framework for assessing mRNA integrity (i.e., presence of full-length or nearly full-length RNAs in a given LNP), we implemented a two-color kinetic fingerprinting assay to simultaneously detect two distinct, well-separated sequences on the same mRNA. We designed an end-region probe for mRNA1 (nt 2033–2040), expected to be less structurally constrained than the internal site (Fig. 5g). End-region probing produced thousands of accepted traces per FOV with robust repetitive probe binding and dissociation (Fig. 5g (ii)), and two-color experiments enabled simultaneous internal- and end-region interrogation to provide a colocalization-based integrity readout within the same LNP population (Fig. 5h and Extended Data Fig. 5a, b). End-region probing showed higher apparent accessibility than middle-region probing, with a shorter median unbound dwell time (τ_off,med_) and a higher total number of binding and dissociation events (*N*_b+d_) (Fig. 5i and Extended Data Fig. 5c), and dye-swap controls preserved this trend (Extended Data Fig. 5a, b). Fano factor analysis placed end-region behavior between the bursty middle-region signal (without tiles) and the Poissonian tiled condition (Extended Data Fig. 5d). Together, these results establish sequence-specific detection of encapsulated mRNA, resolve mixed-cargo formulations, and provide a site-resolved colocalization framework that could be further developed for assessing mRNA integrity within individual, morphology-preserved LNPs.

## Synergized LNP morphology and single-molecule enumeration of encapsulated mRNA

LNP quality control would benefit from combined information on particle morphology and cargo loading, because size and shape distributions can remain stable even when payload heterogeneity varies widely^22–28^. We therefore integrated orthogonal PINPOINT modules within a single assay framework, combining Nile Red PAINT nanoscopy for LNP size and shape with RNA fluorescence for cargo enumeration (Fig. 6a), to measure batch heterogeneity directly with single-particle resolution. We first estimated mRNA copy number per LNP using single-particle RiboGreen staining intensity, reporting these values as fluorescence-calibrated copy-number estimates. Under matched imaging settings, bare mRNA1 provided a reference intensity distribution for one mRNA copy (Fig. 6b)^20,23,24,27^. At the level of individual surface-captured LNP-associated puncta, encapsulated mRNA1 signals in LNP1001 were broader and shifted to higher RiboGreen intensities than the single-mRNA reference, consistent with a mixture of particles containing one or more mRNA copies (Fig. 6b(iii)). We calibrated encapsulated intensity distributions against the bare-mRNA reference to estimate per-particle copy-number distributions (Fig. 6b(iv)). This calibration is justified by our prior bulk measurements indicating that RiboGreen staining intensity of mRNAs within LNPs permeabilized by 0.008% TX-100 is nearly identical to that of fully liberated mRNAs (Fig. 3b). Across LNP formulations, PINPOINT estimated an apparent mean of 2.0–2.3 RiboGreen-accessible mRNA copies among RiboGreen-positive particles, consistent with low average accessible mRNA content under the assumption that LNP-associated RiboGreen signal arises from encapsulated mRNA made probe-accessible after washing and permeabilization (Fig. 6b (v) and Supplementary Table 5)^23,24^.

**Fig. 6.**
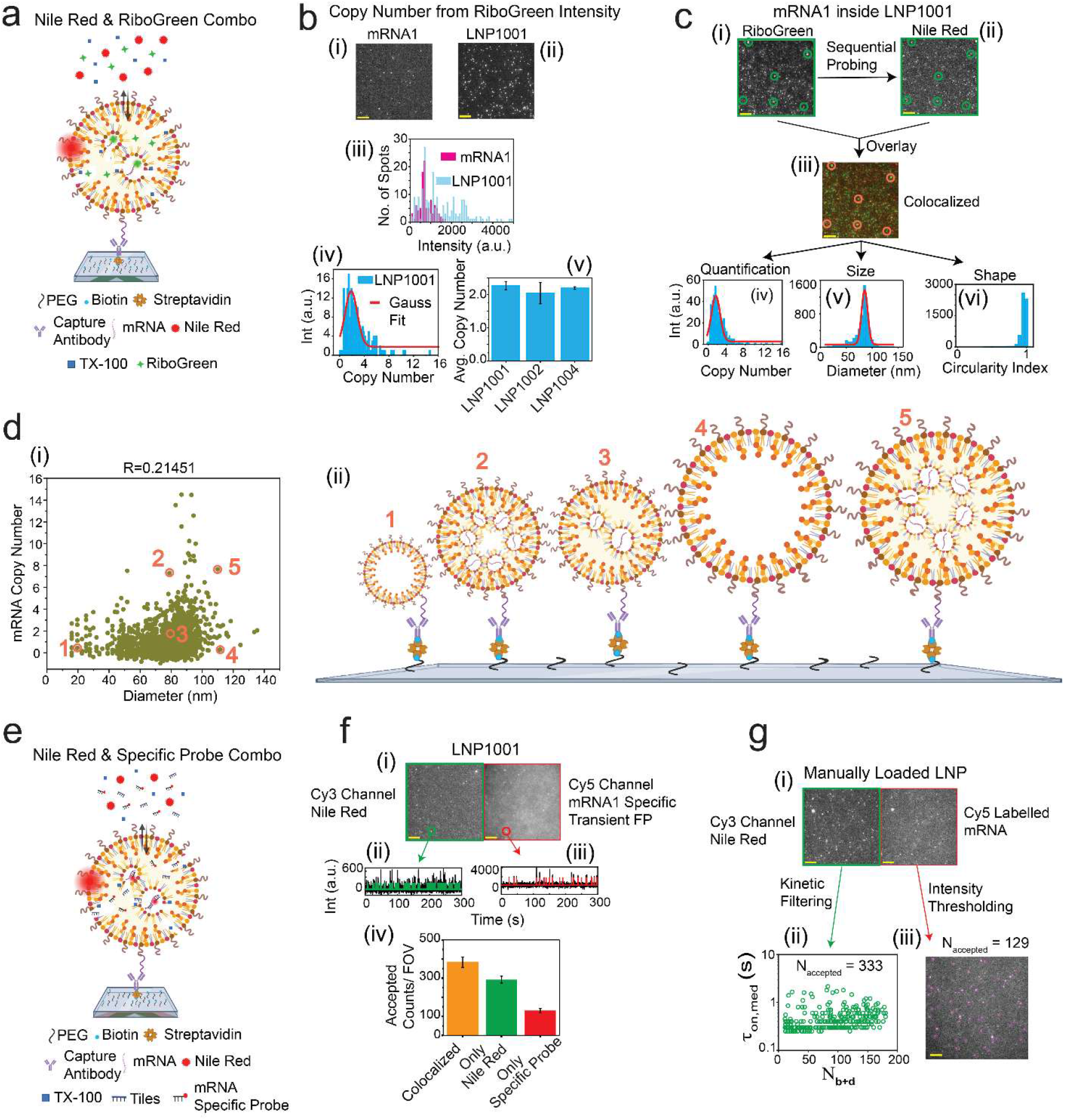
Synergized single-molecule PINPOINT assays enable enumeration and multiparametric quality control of mRNA encapsulation within LNPs. **a,** Schematic of a combined nanoscopic fingerprinting of surface-captured LNPs with Nile Red and RiboGreen enables multiparametric estimation of LNP parameters after permeabilization with optimized TX-100 concentration. Detection is performed using TIRF microscopy. **b**, Average movie frames from TIRF FOV (scale bar, 5 µm) showing binding of RiboGreen to (i) bare mRNA1 and (ii) encapsulated mRNA1 in surface-captured LNP1001. (iii) Comparison of RiboGreen intensity distributions for bare single mRNA1 and individual LNP1001-associated RiboGreen puncta. (iv) Fluorescence-calibrated distribution and Gaussian fit of estimated mRNA copy number per LNP1001 based on RiboGreen intensity. (v) Average fluorescence-calibrated mRNA copy-number estimates for LNP1001, LNP1002, and LNP1004. **c**, Sequential RiboGreen (i) and Nile Red (ii) imaging for encapsulated mRNA1 inside LNP1001, with spatial overlay (iii) enabling colocalization analysis. Multiparametric quantifications from the combined assay: (iv) mRNA copy number; (v) LNP size; (vi) LNP shape (circularity index). Target: LNP1001 at 100 pM; Probe: Nile Red at 1 nM; RiboGreen at 0.5%; DNA tiles at 6 nM; TX-100: 0.008%. **d**, Particle-to-particle heterogeneity in loading relative to size. **(i)** Scatter plot of estimated mRNA copy number (RiboGreen) versus LNP diameter (Nile Red) for individual LNPs (Pearson correlation coefficient shown). Numbered points indicate representative particles. **(ii)** Schematic depictions of the representative particles in d(i), illustrating variability in cargo content at similar particle sizes. **e,** Schematic of a combined PINPOINT assay using Nile Red and mRNA-specific probe enables multiparametric estimation of encapsulated mRNA quality after permeabilization with optimized TX-100 concentration. **f**, Two-channel colocalization experiment for encapsulated mRNA1 detection in LNP1001: (i) Cy3 channel imaging of Nile Red-labeled LNPs and Cy5 channel probing with an mRNA1-specific probe; (ii) and (iii) representative kinetic fingerprinting traces from each channel; (iv) accepted counts per FOV for Nile Red-detected LNPs (deep green), mRNA1-specific probe trajectories (red), and colocalized LNP/probe events (orange), with DNA tiles. Black error bars indicate standard deviation from three independent measurements. Target: LNP1001 at 100 pM; Probe: Nile Red at 1 nM; mRNA1 FP (650–658) _Middle Region_9 nt_Cy5 at 50 nM; DNA tiles at 6 nM; TX-100: 0.008%. **g**, Two-channel experiment for manually loaded Cypridina luciferase (CLuc) mRNA in LNP-ALC0315: (i) Cy3 channel imaging of Nile Red-labeled LNPs and Cy5 channel detection of Cy5-labelled mRNA static spots. (ii) scatterplot of τon,med vs Nb+d for Nile Red-based LNP detection, and (iii) simultaneous counting of Cy5-tagged static spots for estimation of the apparent cargo-positive fraction.

To connect payload to morphology in the same population, we sequentially imaged RiboGreen and Nile Red in the same field of view and performed colocalization analysis (Fig. 6c and Extended Data Fig. 6a). Approximately 60% of Nile Red-detected LNPs colocalized with RiboGreen signal under the optimized permeabilization condition (Extended Data Fig. 6b), implying that ∼40% of particles lacked detectable RiboGreen-accessible mRNA in this assay. This fraction is consistent with the expected RiboGreen-negative population under an approximate Poisson-loading model with a mean near one detectable mRNA per particle. At the particle level, plotting estimated copy number against Nile Red-derived diameter revealed weak (Pearson correlation coefficient R= 0.214) overall coupling between size and cargo content (Fig. 6d), with representative particles illustrating pronounced variability in mRNA copy number among similarly sized LNPs (Fig. 6d(ii)).^20^

We next replaced static RiboGreen readout with kinetic fingerprinting to quantify the particle-level fraction of LNPs containing sequence-detectable cargo in a more sequence-specific manner by assessing colocalization of Nile Red and mRNA-specific probe binding (Fig. 6e and Extended Data Fig. 6c, d). For LNP1001, this approach yielded an apparent cargo-positive fraction of ∼60%, based on the fraction of Nile Red-detected particles that also produced accepted mRNA1-probe trajectories (Fig. 6f). For a subset of mRNA1-positive LNPs, the mRNA-specific probe trajectories exhibited discrete multistep intensity levels, consistent with simultaneous binding of multiple probes to multiple accessible mRNA copies within the same particle (Supplementary Fig. 7). We therefore interpret these traces as qualitative support for multi-copy loading and as a potential trajectory-based route to cargo enumeration, rather than as the primary copy-number measurement. Finally, in an independently prepared formulation in which Cypridina luciferase (CLuc) mRNA was manually loaded into LNP-ALC0315, the same colocalization logic yielded a lower apparent cargo-positive fraction of 33% (Fig. 6g), consistent with the expected variability of manual loading^19^. Together, these synergized assays quantify copy-number distributions, cargo-positive fractions and morphology on the same single-LNP populations, enabling multiparametric batch fingerprinting for formulation and process optimization.

## Conclusion

Lipid nanoparticles have enabled the rapid clinical translation of mRNA medicines, but their multicomponent, self-assembled nature makes them heterogeneous and sensitive to formulation scale up, manufacturing, and handling idiosyncrasies^29,30^. Because routine assays typically only report population averages or require disruptive processing^19,20^, ensuring batch consistency, linking particle structure to payload, and understanding attributes that influence efficacy and in vivo fate remain challenging. Here we introduced PINPOINT, a single particle nanoscopy platform that quantifies multiple critical quality attributes of mRNA–LNPs in a unified, operationally non-destructive workflow. Under the optimized sub-CMC TX-100 condition, LNPs retain Nile Red-detectable morphology rather than dissolving, while becoming sufficiently permeable to permit repeated access to encapsulated mRNA. PINPOINT provides multiparametric characterization of individual particles across particle size, circularity-based shape metrics, cargo identity, cargo accessibility, and loading. Super-resolution Nile Red imaging resolves LNP size and shape distributions and detects handling-induced changes such as emergent non-circular subpopulations after freeze–thaw stress, providing a potentially useful quality control measure in the event of variable sample handling or storage.

A controlled sub-CMC TX-100 permeabilization window enables probe access to encapsulated RNA while preserving particle morphology and reveals that cargo accessibility is dynamic and can be sustained by surfactant replenishment. Sequence-specific kinetic fingerprinting achieves high discrimination even for closely related mRNAs (∼80% sequence identity in the probed region and 98% identity across the full-length sequence), and targeted DNA tiling regularizes structure-gated binding to improve quantification. Multi-region probing provides site-resolved information about RNA accessibility and a potential colocalization-based framework for assessing integrity within individual LNPs, and combined assays integrating Nile Red with RiboGreen or mRNA-specific probes estimate accessible mRNA copy-number distributions and particle-level cargo-positive fractions, exposing heterogeneous loading and co-detection patterns that are obscured by ensemble averaging ^22–28^. In the multiplexed detection study, cargo identity and loading metrics provided context beyond morphology-only signatures and distinguished the defined cargo compositions from an independent cargo-negative control. More broadly, PINPOINT reframes particle-to-particle variability from a confounder into a quantitative descriptor that can be tracked across batches, formulation changes and storage conditions, with direct relevance to quality control and rational design.

PINPOINT has limitations. The current implementation is not optimized for high-throughput screening or in-line manufacturing readout; present throughput is on the order of 10^4^ particles per hour, and the current manual workflow requires approximately 3 h from sample preparation to result. PINPOINT also requires dilution of LNP samples to low-picomolar working concentrations to achieve spatially separated single-particle capture, so potential dilution-dependent effects should be controlled by matched handling conditions and reference formulations. Analysis also requires specialized software and careful parameterization for trace extraction, kinetic filtering and colocalization^31,32^. Detection depends on probe accessibility and background conditions, and permeabilization introduces time-dependent accessibility that must be controlled with care to limit false negatives due to re-sealing of membranes. The tiling strategy improves local access but adds an assembly step that may not be desirable in all settings^38^, and whose efficacy may be reduced by highly structured cargo RNAs. Throughput, speed and robustness could be improved using pre-functionalized imaging surfaces, faster capture chemistries, fluorogenic probes for faster imaging^45,46^, automated microfluidic buffer exchange for continuous introduction of TX-100, and greater standardization of calibration using reference materials. Analysis could also be further automated using machine-learning models trained on labels derived from hidden Markov modeling (HMM) and thresholding. Such models could reduce manual parameter tuning and improve resilience across conditions by classifying trajectories (specific versus nonspecific, high versus low confidence), flagging outliers, and automating particle morphology and colocalization calls from reconstructed features ^47^. Expanded probe panels and spectral demixing or kinetic encoding could further increase multiplexing without additional labelling of LNP components^32^. Overall, PINPOINT provides an integrated, amplification-free strategy for quantitative, sequence-resolved interrogation of individual mRNA–LNPs, enabling resolution of heterogeneity inaccessible to bulk methods to support batch consistency and optimization of RNA therapeutic delivery systems.

## Methods

### Materials

All DNA and RNA oligonucleotides were purchased from Integrated DNA Technologies (IDT) with high-performance liquid chromatography purification. Oligonucleotide sequences are listed in Supplementary Table 1. Unless otherwise noted, oligonucleotides were dissolved in TE buffer (10 mM Tris-HCl, 0.1 mM EDTA, pH 8.0) to 100 µM, aliquoted and stored at −20 °C (DNA) or −80 °C (RNA). mRNA1, mRNA2, and LNP formulations LNP1001, LNP1002, and LNP1004 were provided as research materials; their cargo compositions and sample concentrations are summarized in Supplementary Table 2. LNP-ALC0315 was obtained from Cayman Chemical. Cypridina luciferase (CLuc) mRNA was synthesized in vitro using the HiScribe T7 mRNA with CleanCap Reagent AG kit (New England Biolabs, E2080S) according to the manufacturer’s instructions. Cy5-labelled mRNA was encapsulated at 50 ng µl⁻¹ using the LipidLaunch ALC0315 LNP Kit (Loadable) (Cayman Chemical, 702750). Monoclonal anti-PEG capture antibody (Abcam, ab170969) was used for LNP immobilization. Capture antibodies were bovine serum albumin (BSA)-free and azide-free to enable biotinylation using NHS–biotin reagents (where applicable). Nile Red (CAS 7385-67-3) and Triton X-100 (TX-100) (CAS X100-5ML) were purchased from Sigma-Aldrich (MilliporeSigma). The Quant-iT RiboGreen RNA Assay Kit and RiboGreen RNA reagent were purchased from Invitrogen (Thermo Fisher Scientific, R11490). All additional chemicals used for buffer preparation were obtained from Sigma-Aldrich or Thermo Fisher Scientific. DNA LoBind tubes (Eppendorf) were used for all experiments to minimize sample adsorption.

### Biotinylation of capture antibody

Monoclonal antibodies were biotinylated by reaction with biotin *N*-hydroxysuccinimidyl ester (Sigma-Aldrich, Catalog H1759−100), an amine-reactive NHS ester, in a solution containing a of 5:1 molar ratio of biotin: antibody, and incubated at room temperature for 1h in 1× PBS at pH 7.4 as described in previous studies^32,43^. The biotin-IgG conjugates were purified through Zeba Spin desalting columns (Thermo Fisher, Catalog 89882, 7K MWCO) according to the manufacturer’s protocol and underwent overnight dialysis (Slide-A-Lyzer Dialysis Cassette, Thermo Fisher, 3.5K MWCO) at 4 °C against 1× PBS, pH 7.4. Biotinylation was estimated with an electrophoretic mobility shift assay with or without excess streptavidin and ranged between 70% and 80%. The final capture antibodies were aliquoted and stored at −80 °C.

### Cy5-labelled mRNA synthesis

Cypridina luciferase (CLuc) mRNA was synthesized *in vitro* utilizing the HiScribe® T7 mRNA with CleanCap® Reagent AG kit (New England Biolabs, E2080S), in accordance with the manufacturer’s instructions. The kit-supplied linearized DNA template was used. To enable fluorescent labelling, 30% of the uridine triphosphate (UTP) was substituted with Cy5-UTP (Apexbio Technology LLC, B8333) during transcription. Following synthesis, the mRNA was purified with the MEGAclear™ Transcription Clean-Up Kit (Thermo Fisher Scientific, AM1908), again according to the manufacturer’s protocol.

### Encapsulation of mRNA in LNP-ALC0315

Cy5-labelled CLuc-mRNA was encapsulated at a concentration of 50 ng/μL using the LipidLaunch™ ALC0315 LNP Kit (Loadable) (Cayman Chemical, 702750), following the provided protocol. After mRNA loading, the size distribution of the resultant LNPs was determined by dynamic light scattering (DLS) using the Dynapro Nanostar instrument (Wyatt Technology).

### RiboGreen assay in steady state fluorescence

Steady-state fluorescence spectra were acquired on a Horiba Scientific fluorometer (Fluoromax Plus-C) using quartz cuvettes. Blank spectra were collected in 1× PBS (pH 7.4; Gibco). For TX-100 titrations, samples (final volume 200 µl) were prepared in 1× PBS (pH 7.4) containing LNPs (100 pM), RiboGreen reagent (1:200 dilution; Thermo Fisher Scientific) and Triton X-100 (TX-100; 0–1% v/v, as indicated). Samples were excited at 480 nm and emission was recorded from 490 to 700 nm using 3 nm excitation and emission slit widths. Instrument signal and reference corrections were enabled (S1C and R1C), and spectra are reported as the ratio of sample detector to reference detector (S1C/R1C).

### Dynamic Light Scattering

Hydrodynamic diameter distributions of LNPs were measured by dynamic light scattering (DLS) using a DynaPro NanoStar (WDPN-10; Wyatt Technology). Samples were prepared at 100 pM in PBS (pH 7.4), and 10 µl of each sample was loaded for measurement. Data were collected and analysed using DYNAMICS software (Wyatt Technology). Particle diameters were determined by instrument software using cumulant analysis. For reporting an average diameter from the diameter histogram, we calculated the amplitude-weighted centroid using equation 1,

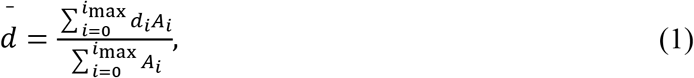

where *d_i_* is the diameter at bin *i* and *A_i_* is the corresponding histogram amplitude.

### Preparation of glass slide surfaces for single-molecule microscopy

Glass coverslips (No. 1.5, 24 × 50 mm; VWR, 48393-241) were PEG-passivated using a 1:10 molar mixture of biotin–PEG–SVA and mPEG–SVA (Laysan Bio; Biotin-PEG-SVA-5K-100MG and MPEG-SVA-5000-1g) following a standard protocol^31,32^. After PEGylation, coverslips were stored in a Coplin jar wrapped with aluminum foil in a nitrogen-purged cabinet and used within 4 weeks. For imaging experiments, multiwell chambers were constructed by cutting micropipette tips (Thermo Fisher Scientific, 02-682-261) approximately 1 cm from the wide end, discarding the narrow portion and placing the wide end in contact with the PEGylated surface to define a well. Wells were sealed to the coverslip using epoxy (Ellsworth Adhesives, 4001).

### Single-molecule fluorescence microscopy

All single-molecule experiments were performed using TIRF microscopy on an Oxford Nanoimager (ONI) (refer to https://oni.bio/nanoimager/ for camera, illumination, and objective specifications; this unit has a 532 nm excitation line in place of the standard 561 nm line). The ONI uses a 100 x 1.4NA oil-immersion objective and Z-lock control setting for autofocus. To prevent overheating, two cooling plates were added to the ONI externally, secured with metal clamps, to allow cold water from a water bath to flow through them since the ONI’s built in temperature control system could not maintain imaging temperatures below 25 °C. Before imaging, we set the ONI to the desired temperature and adjust the water bath temperature accordingly after laser activation to stabilize imaging temperature at 25 °C at our ambient temperature of ∼22 °C. An illumination angle of approximately 54° was set for TIRF. Samples were excited at 532 nm (at 30% laser power) and 640 nm (at 35% laser power) to capture Cy3 and Cy5 fluorescence emission, respectively, at an optimal signal-to-noise-ratio (S/N). Each FOV was observed for 5 min at a signal integration time of 100 ms per frame unless stated otherwise. Experiments that require colocalization were recorded simultaneously with two laser channels active.

### Fluorescent probe selection & imaging solution

Probe design was guided by the requirement for reversible target binding to enable repeated association–dissociation cycles for kinetic fingerprinting. For sequence-specific oligonucleotide probes, we targeted melting temperatures near room temperature (20–25 °C) and used short probe lengths (typically 8–10 nt; 11 nt where noted) to achieve dissociation kinetics on a timescale of ∼1 second. Probe GC content was kept ≤50% when possible. At least 2 nucleotides were left unpaired between adjacent FP and CP/tile binding sites to reduce potential stacking interactions that would influence binding kinetics. Probe and tile sequences are provided in Supplementary Table 1. Unless stated otherwise, single-molecule imaging was performed in 2× PBS (pH 7.4; Gibco) supplemented with an oxygen scavenging system consisting of 5 mM 3,4-dihydroxybenzoic acid (Fisher Scientific, AC114891000), 0.05 mg ml⁻¹ protocatechuate 3,4-dioxygenase (Sigma-Aldrich, P8279-25UN), and 1 mM Trolox (Fisher Scientific, 218940050). Where indicated, imaging buffers additionally contain DNA tiles (6 nM each) and/or TX-100 (0.008% v/v; freshly prepared from a 2% stock) and fluorophore-labelled oligonucleotide probes (typically 50 nM).

For the Nile Red assay, Nile Red was serially diluted in DMSO and then diluted into 2× PBS immediately before imaging to a final concentration of 1 nM. For the RiboGreen assay, imaging buffer contained 2× PBS (pH 7.4), RiboGreen reagent from the Quant-iT™ RiboGreen RNA Assay Kit (0.5% v/v, corresponding to a 1:200 dilution of the supplied reagent stock) and TX-100 (0.008% v/v).

### Capturing strategy & kinetic fingerprinting assay

All sample preparation was performed in low-adhesion 1.7 ml microcentrifuge tubes (GeneMate). Unless otherwise noted, all incubations were carried out at room temperature. For surface preparation, each sample well was washed with 100 µl of T50 buffer (10 mM Tris-HCl, 50 mM NaCl, 1 mM EDTA, pH 8.0) for 5 min. After removal of T50, streptavidin (40 µl, 1 mg ml⁻¹) was added and incubated for 10 min. Excess streptavidin was removed and wells were washed three times with 100 µl of 1× PBS.

For LNP assays, wells were incubated with biotinylated anti-PEG capture antibody (40 µl, 100 nM) for 30 min. Excess antibody was removed and wells were washed three times with 100 µl of 2× PBS. LNP stocks (0.1 µM) were diluted 1,000-fold in 2× PBS to a working concentration of 100 pM (unless otherwise specified) and added to the well. LNP solutions were handled by gentle pipetting and were not vortexed. After a 1h capture incubation, excess LNPs were removed and wells were washed three times with 100 µl of 2× PBS. Imaging solution (100 µl; composition described above) was then added and samples were imaged promptly by TIRF microscopy.

For bare mRNA assays, wells were pre-incubated with a biotinylated 3′ DNA capture probe (100 µl, 100 nM) for 10 min. Excess capture probe was removed and wells were washed three times with 100 µl of 2× PBS before adding sample. For tiled mRNA experiments, unformulated mRNA was mixed with DNA tiles (final mRNA concentration 10 pM; tile concentration as indicated) and annealed using a thermocycler program of 95 °C for 2 min, 40 °C for 8 min, followed by cooling in a room-temperature water bath for 5 min. The annealed sample was then added to the well and incubated for 1 h to capture tiled mRNA on the surface. Excess mRNA was removed, wells were washed three times with 100 µl of 2× PBS, and 100 µl of imaging solution was added immediately before TIRF imaging (Supplementary Fig. 3).

### Analysis of single-molecule data

PINPOINT single-molecule kinetic fingerprinting datasets were analyzed using custom MATLAB scripts to quantify probe-binding kinetics and perform digital counting, following established diffraction-limited kinetic fingerprinting workflows (Supplementary Fig. 4)^31,32^. Candidate regions of interest (ROIs) were identified by computing an intensity fluctuation map, defined as the average absolute frame-to-frame change in intensity at each pixel across the movie. Local maxima in the fluctuation map were used to seed ROIs, which were defined as 3 × 3-pixel regions centred on each maximum. For each ROI, background-subtracted integrated intensity was computed for every frame to generate an intensity-versus-time trajectory.

Trajectories were idealized using hidden Markov modelling (HMM) with a custom implementation of vbFRET. Idealized state sequences were used to extract kinetic fingerprint parameters, including *N_b+d_* (number of binding and dissociation events), τ_on,med_ and τ_off,med_ (median dwell times in bound and unbound states), and τ_off,max_ (maximum unbound dwell time). Signal-to-noise ratio (S/N) was calculated as the standard deviation of the fluorescence intensity divided by the mean intensity difference between the bound and unbound states. For each probe modality, thresholds for intensity, S/N and kinetic parameters were optimized using an automated routine within the software to maximize the trajectories classified as accepted (positive detection events) in positive controls while minimizing accepted counts in corresponding negative controls; optimized parameter sets are shown in Supplementary Table 4.

Colocalization analysis (for sequential or dual-colour assays) was performed in MATLAB using ROI coordinate information retained during movie processing and reconstruction, enabling registration of spots across channels and/or acquisitions and enumeration of colocalized events.

### Fitting of the cumulative frequency of dwell times

Dwell-time distributions were analyzed in MATLAB using cumulative frequency (cumulative distribution) fitting. For each dataset, bound- and unbound-state dwell times extracted from HMM-idealized trajectories were pooled and binned at the camera exposure time (100 ms). The cumulative frequency *y*(*x*) as a function of dwell time *x* was then fit with a single-exponential model,

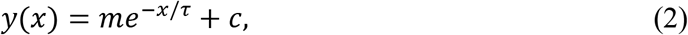

where *m*, τ and *c* are fit parameters. For cumulative distributions normalized to span from 1 to 0, *m* was constrained to −1, *c* corresponds to the long-time offset, and τ represents the characteristic (mean) dwell time of the underlying exponential process. Fits were accepted when the sum of squared errors was < 0.05 and *R*^2^ was > 0.99, indicating that a single-exponential model adequately described the cumulative distribution and that τ provided a representative dwell-time estimate.

### Super-resolution analysis (Nile Red)

Super-resolution LNP images were reconstructed from the fluctuating fluorescence signals arising from repeated, transient association of single Nile Red molecules with hydrophobic LNP domains and/or intrinsic blinking of bound dyes (Fig. 2) according to the PAINT method^34^. Individual binding events were localized in each frame by least-squares fitting to a 2D Gaussian function, yielding per-event centroid coordinates which were corrected for translational stage drift (generally ≤ 1 pixel) during the measurement using subpixel registration with the discrete Fourier transform cross-correlation method^34,35,48^. Each localization was then represented as a smaller Gaussian at the centroid coordinates with a standard deviation equal to the average localization error (∼30 nm). Gaussian representations for localizations from all frames were then pooled to generate a super-resolution reconstruction for all particles in the field of view, which was used to estimate particle diameter and area (defined by a boundary threshold of ½ maximal localization density within each particle, akin to a full-width at half maximum metric of size), and to compute a circularity index (range 0-1, where 1 = maximum circularity) using the MATLAB regionprops command. The circularity index is equal to

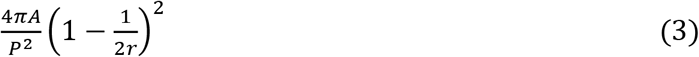

where *A* is the area of the bounded region, *P* is the perimeter of the bounded region, and 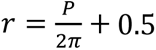. Analyses were restricted to particles containing regions with at least two spatially overlapping localizations (i.e., within localization error of one another) to filter out extraneous/nonspecific events. Size distributions and summary statistics were then compiled across fields of view for comparisons between batches and conditions (Fig. 2 and Supplementary Table 3).

### Burst analysis and Fano factor calculations

Bursting in probe-binding trajectories were quantified using a modified Rank Surprise (RS) method adapted from Gourévitch et al^44^. Interspike intervals (ISIs) were defined as the time between consecutive probe-binding events for each accepted single-molecule trajectory (from HMM-idealized traces). For burst calling, we implemented a global RS approach in MATLAB in which ISIs from all trajectories within an experimental condition were ranked jointly, so that burst identification was not biased by molecules with unusually high or low event counts. The RS algorithm was then used to identify contiguous sequences of short ISIs (“bursts”) and to assign burst start and end points, which were mapped back to each trajectory to preserve single-molecule burst profiles. RS parameters were set to a maximum ISI of 40 s for events to be considered within a burst and an RS cutoff (α) of 3.

Fano factor analysis was used to test whether spike trains deviated from Poisson statistics. For each condition, the Fano factor was calculated as the variance of spike counts divided by the mean spike count within a time window *T*. For each trajectory, a window of length *T* was randomly sampled 100 times (independent random seeds) and Fano factors were averaged across samplings and then across trajectories. Calculations were performed for *T* = 5,10,20,40 and 60 s (as plotted). Poisson reference behaviour was generated by simulating spike counts using MATLAB’s Poisson random number generator with a mean rate matched to the experimentally observed average burst ISI and using the same number of samplings. The 95% confidence interval for Poisson behaviour was computed using gamma-distribution bounds, gam*i*nv([0.025,0.975], (*n* − 1)/2,2/(*n* − 1)), where *n* is the sample size.

### mRNA copy number estimation

mRNA copy number per LNP was estimated from single-particle RiboGreen fluorescence by normalizing encapsulated-particle intensities to single-mRNA reference^23^. Bare mRNA1 was imaged under the same optical settings used for LNP measurements to obtain a one-copy intensity reference. The bare-mRNA intensity histogram fitted with a Gaussian model, and the fitted mean was taken as the characteristic intensity of one mRNA copy. Encapsulated mRNA intensity distributions were obtained by imaging RiboGreen-accessible RNA in permeabilized, surface-captured LNPs. For each LNP, the estimated copy number was calculated as the particle intensity divided by the one-copy reference intensity, and batch-level copy-number distributions and mean copy numbers were then obtained (Fig. 6b and Supplementary Table 5).

## Data availability

The data supporting the findings of this study are available within this Article and its Supplementary Information. Source data are provided with this paper. Any additional data are available from the corresponding author upon request.

## Code availability

The code used in this study has been described in reference^31–32^ and is available through a DeepBlue repository. Any additional information concerning the code is available from the corresponding author upon request.

## Acknowledgements

We thank Pfizer Inc., specifically Patrick Lim Soo, Chelsea R. Thorn, Anthony Barry, and John Androsavich, for providing the mRNA and LNP samples used in this study. This work was supported by NIH MIRA grant R35 GM131922 (N.G.W.).

## Author contributions

P.B., T.C. and N.G.W. conceived the idea. P.B. and T.C. designed the experiments. P.B., T.C. and N.E.W. performed most experiments. P.B., T.C., N.E.W. and A.J.B. analyzed, discussed the data and created the figures. P.B. wrote the initial draft of the manuscript. N.G.W. supervised the project. All authors reviewed and edited the manuscript.

## Additional information

**Supplementary information** Supplementary information is available in the online version of the paper.

---

**Extended Data Fig. 1.**
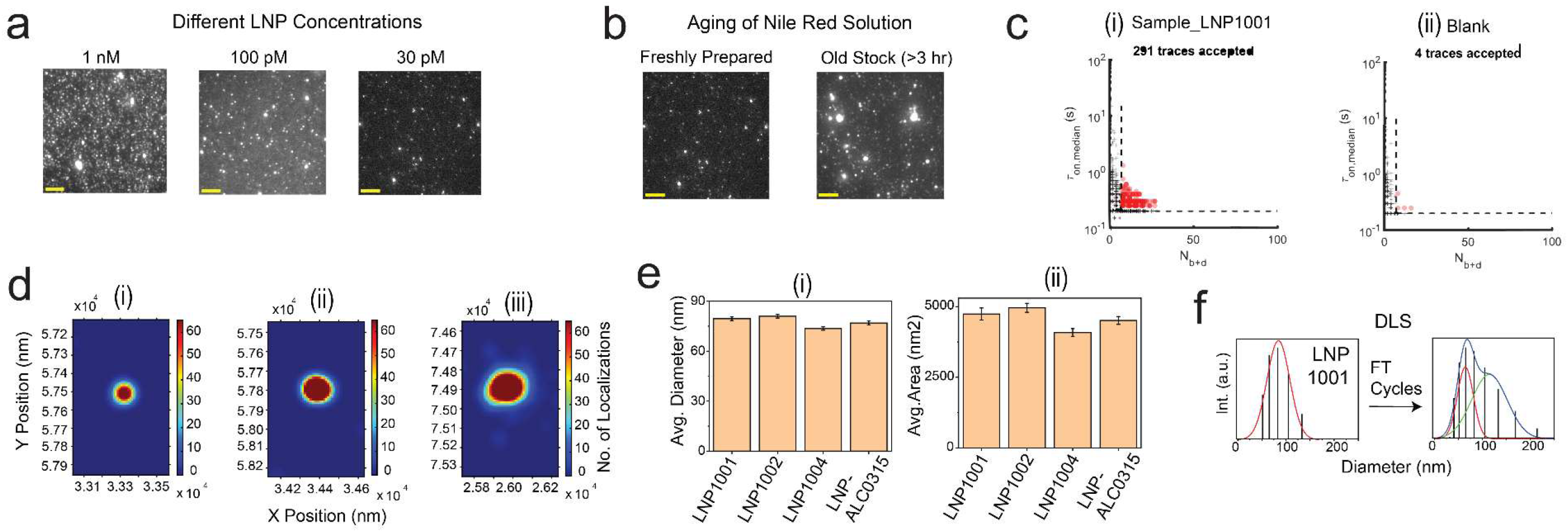
Single-molecule detection and spatial characterization of LNPs. **a,** Impact of different LNP concentrations (1 nM, 100 pM, 30 pM) on imaging. **b,** Effect of aging of Nile Red solution (freshly prepared vs. old stock >3 hr) on imaging quality. Sample: LNP1001; Temperature: 25°C; Probe: Nile Red at 1 nM. **c,** Scatterplots of dwell time analysis (i.e., τ_on,med_ versus τ_off,med_) for all intensity-versus-time trajectories observed within a single field of view in the presence (i) or absence (ii) of 100 pM LNP1001. Dashed lines indicate thresholds (minimum or maximum) for accepting a trajectory as evidence of a single LNP molecule. Points indicated by “+” represent trajectories that do not pass filtering for intensity, signal-to-noise ratio, and/or kinetics, and are not considered sufficient evidence to detect LNP1001. Points indicated by red-filled circles represent trajectories that pass filtering and are considered positive detection events of single LNP molecules. **d**, Super-resolution images of representative small (i), medium (ii), and large (iii) LNPs in the same FOV. Colour scale represents the number of localization events; x and y axes are spatial coordinates (nm) and are identical across panels. **e**, Average diameter (i) and area (ii) of LNP1001, LNP1002, LNP1004, and LNP-ALC0315 determined from super-resolution analysis using Nile Red PINPOINT assay. Data are presented as mean ± SD. Error bars indicate the standard deviation of three independent measurements. Temperature: 25°C; Acquisition time: 5 min/FOV; Probe: Nile Red at 1 nM. **f,** Average diameter determined from DLS before and after freeze–thaw cycles of LNP1001.

**Extended Data Fig. 2.**
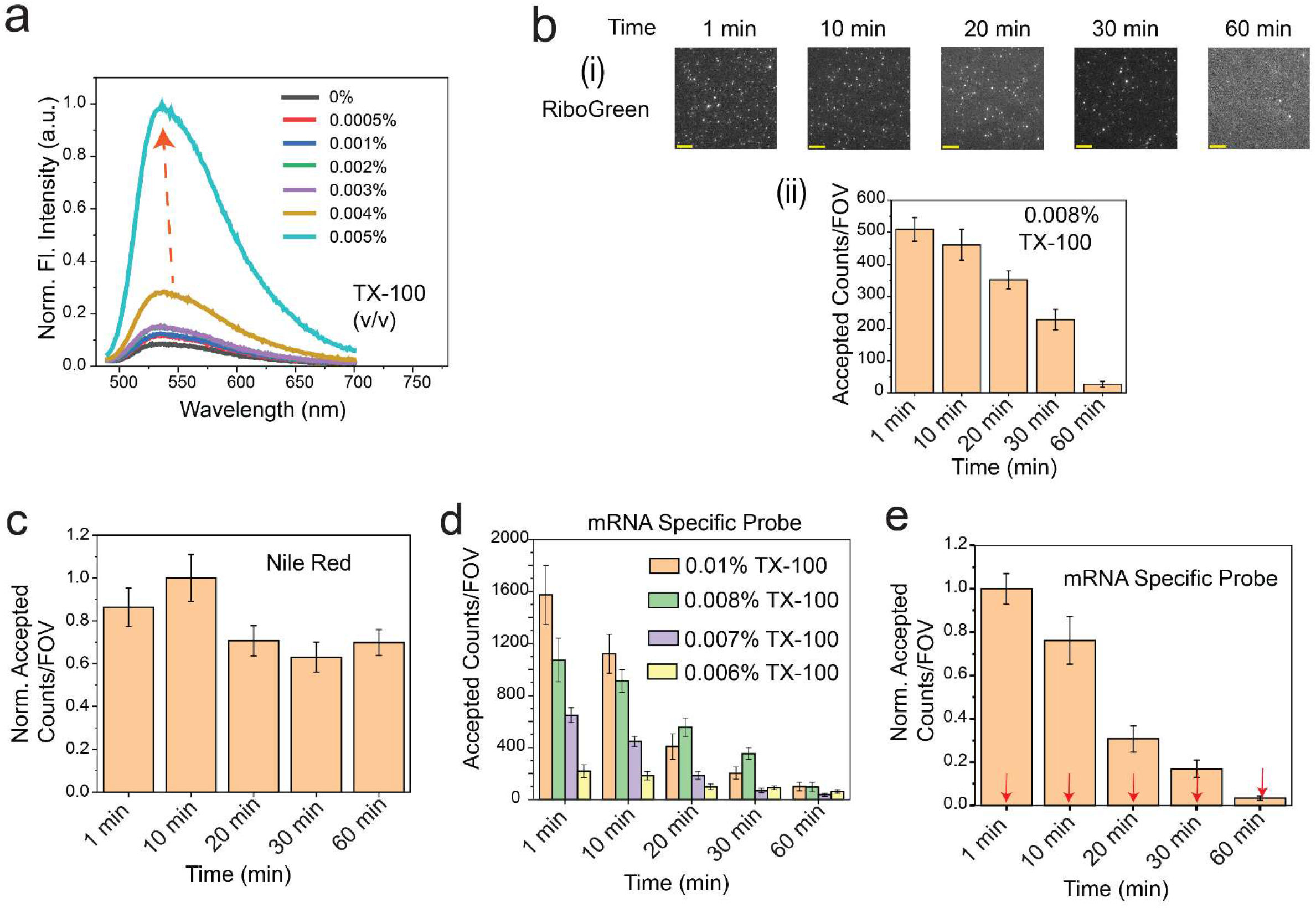
Fluorescence-based assessment of LNP cargo accessibility under varying TX-100 conditions and time. **a,** Steady-state fluorescence spectra (normalized fluorescence intensity vs wavelength) of LNP1001 with RiboGreen at 0.5% across a range of TX-100 concentrations (0%–0.005% v/v), highlighting the sensitivity window for membrane permeabilization. **b**, RiboGreen imaging and quantification of mRNA1 accessibility inside LNP1001 in the presence of 0.008% TX-100. (**i**) Single movie frames from a representative TIRF microscope field of view (FOV; scale bar, 5 µm) show bright puncta corresponding to RiboGreen bound at or near mRNA1 inside LNP1001. (**ii**) Quantification of accepted counts per FOV as a function of time (1, 10, 20, 30, and 60 min). Temperature: 25 °C; Acquisition time: 15 s/FOV; Probe: RiboGreen at 0.5%. **c**, Quantification (normalized accepted counts per FOV) of LNP particles detected over time in the same system by Nile Red super-resolution imaging. Probe: Nile Red at 1 nM. **d**, Quantification of accepted counts per FOV for mRNA1-specific probes inside LNP1001 over time at multiple TX-100 concentrations within the optimized range. Acquisition time: 5 min/FOV; Probe: mRNA1 FP (650–658) _Middle Region_9 nt_Cy5 at 50 nM. **e**, Quantification (normalized accepted counts per FOV) of mRNA1-specific probe accessibility inside LNP1001 over time in the presence of 0.008% TX-100, with external addition of DNA tiles (6 nM) at each time point (indicated by red arrows). Acquisition time: 5 min/FOV; Probe: mRNA1 FP (650–658) _Middle Region_9 nt_Cy5 at 50 nM; DNA tiles at 6 nM. Black error bars represent standard errors of the mean from three independent replicates.

**Extended Data Fig. 3.**
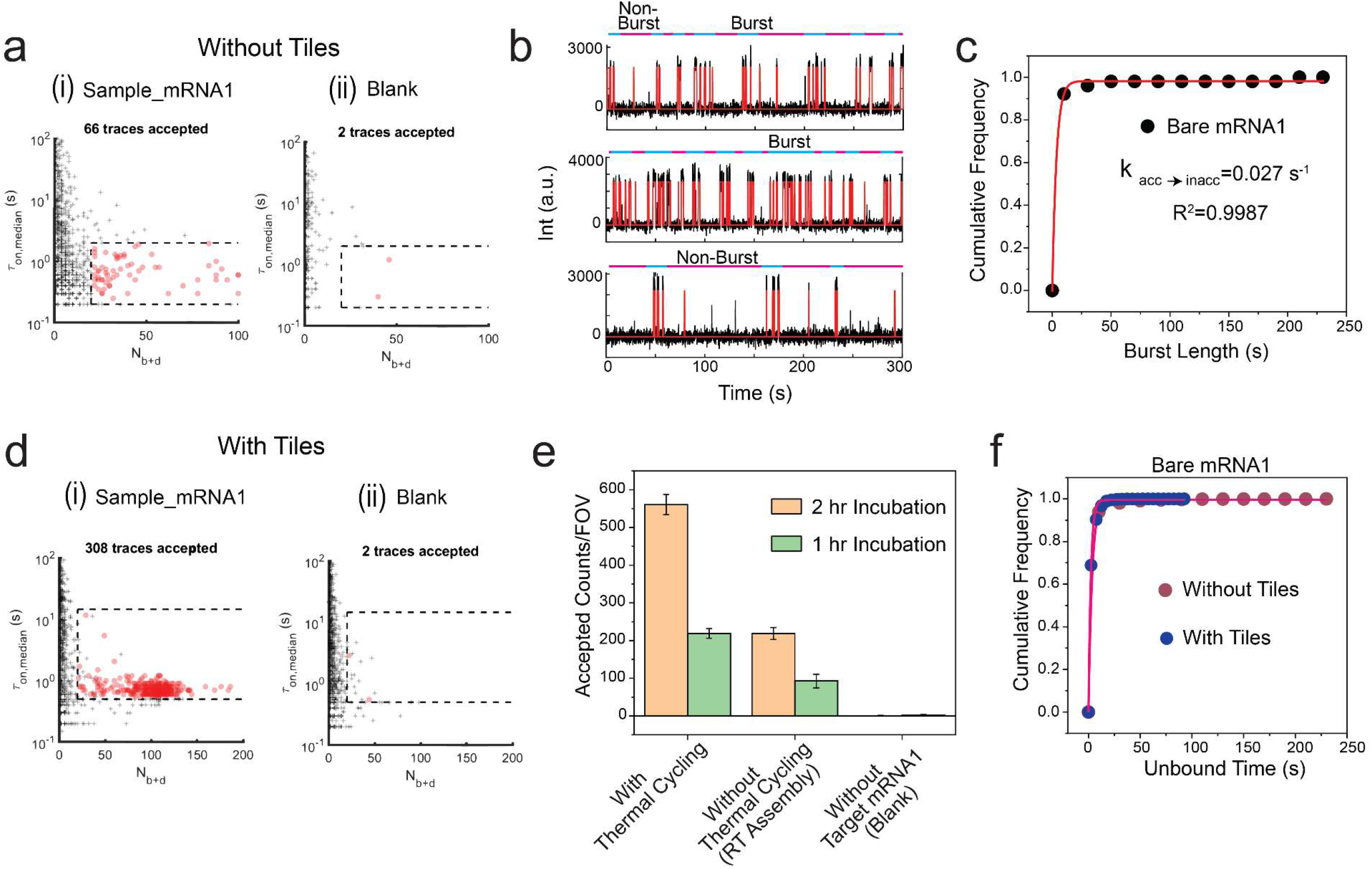
Influence of DNA tile assembly and molecular accessibility on single molecule bare mRNA1 detection. **a,** Scatterplots of dwell time analysis (τ_on,med_ versus τ_off,med_) for all intensity-versus-time trajectories observed within a single field of view in the presence (i) or absence (ii) of 10 pM bare mRNA1 without tiles. Dashed lines indicate thresholds (minimum or maximum) for accepting a trajectory as evidence of a single mRNA1 molecule. **b**, Three representative intensity-versus-time trajectories for the interaction of bare mRNA1 (without tiles) with the sequence-specific probe, showing burst (cyan bars) and non-burst (magenta bars) target binding periods detected through spike train analysis. **c**, Cumulative distribution function of burst duration fitted with a single-exponential equation, yielding the rate constant (k_acc→inacc_) for bare mRNA1 accessibility transitions, reflecting secondary structure effects. **d**, Scatterplots of dwell time analysis (τ_on,med_ versus τ_off,med_) for all intensity-versus-time trajectories observed within a single field of view in the presence (i) or absence (ii) of 10 pM bare mRNA1 with DNA tiles. Dashed lines indicate thresholds (minimum or maximum) for accepting a trajectory as evidence of a single mRNA1 molecule. **e**, Effect of thermal cycling during DNA tile assembly on accepted counts of bare mRNA1 with tiles at two different sample incubation times. Black error bars represent standard errors of the mean from three independent replicates. **f**, Cumulative unbound time histograms comparing bare mRNA1 with and without tiles, with τ_off,med_ values determined from single-exponential fitting of cumulative unbound time distributions of accepted single-molecule traces. Temperature: 25°C, Acquisition time: 5 min/FOV; Probe: mRNA1 FP (650–658) _Middle Region_9 nt_Cy5 at 50 nM; DNA tiles at 6 nM.

**Extended Data Fig. 4.**
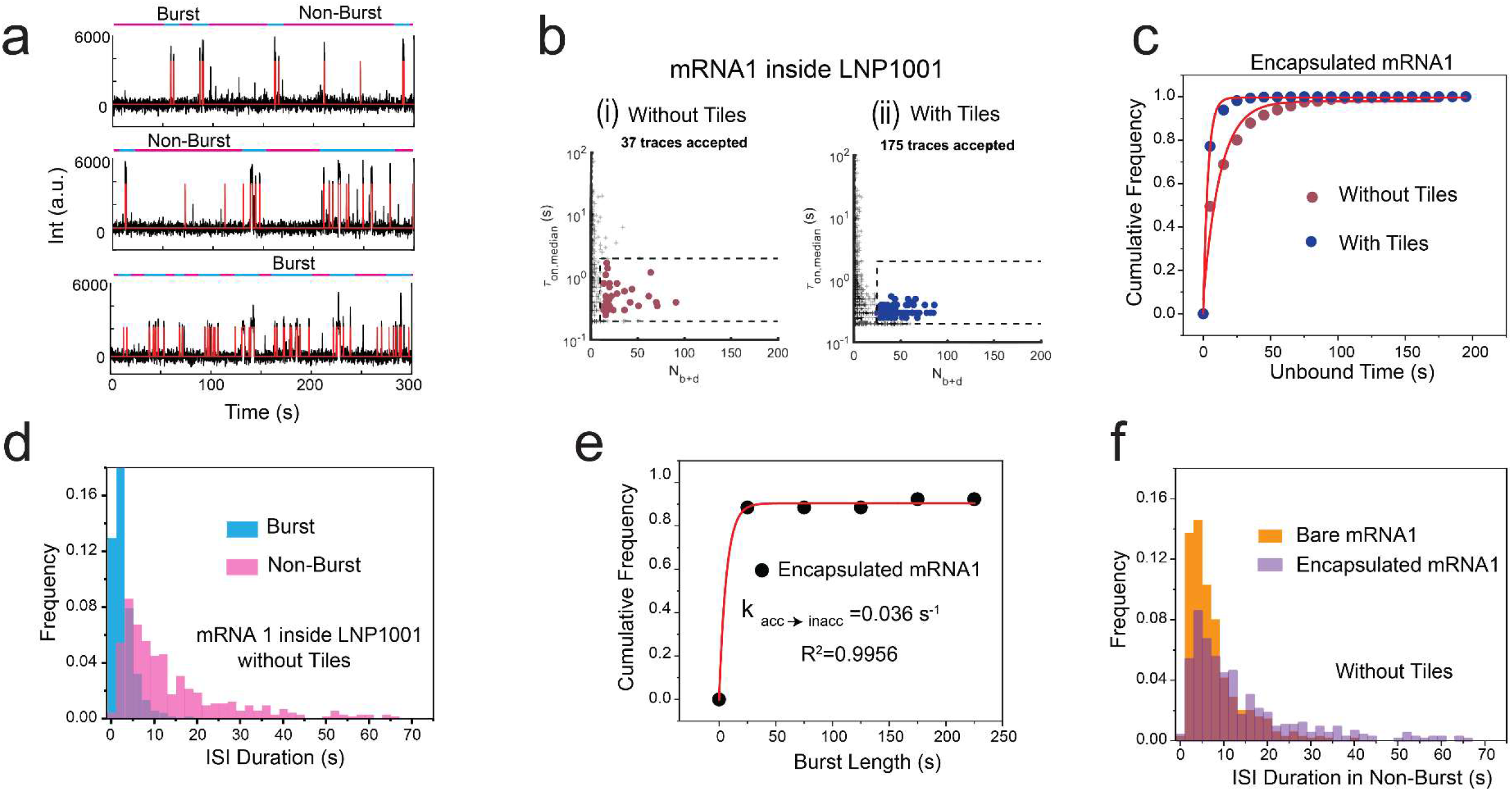
Single-molecule kinetics and accessibility of encapsulated mRNA1 with and without DNA tile enhancement. **a,** Three representative intensity-versus-time trajectories for the interaction of encapsulated mRNA1 (inside LNP1001, without tiles) with the sequence-specific probe, showing burst (cyan bars) and non-burst (magenta bars) target binding periods detected via spike train analysis. **b**, Scatterplots of dwell time analysis (τ_on,med_ versus τ_off,med_) for encapsulated mRNA1 in LNP1001 (i) without tiles and (ii) with tiles. Dashed lines indicate thresholds (minimum or maximum) for accepting a trajectory as evidence of a single mRNA1 molecule. **c**, Cumulative unbound time histograms comparing encapsulated mRNA1 with and without tiles, with τ_off,med_ values determined from single-exponential fitting of cumulative unbound time distributions of accepted single-molecule traces. **d**, Cumulative histogram displaying the distribution of interspike intervals (ISIs) during burst (cyan) and non-burst (magenta) periods for encapsulated mRNA1 without tiles. **e**, Cumulative distribution function of burst duration fitted with a single-exponential equation, yielding the rate constant (k_acc→inacc_) for encapsulated mRNA1 accessibility transitions, reflecting secondary structure effects. **f**, Comparison of ISI duration distributions in non-burst periods for bare mRNA1 versus encapsulated mRNA1 (both without tiles). Temperature: 25°C; Acquisition time: 5 min/FOV; Target: LNP1001 at 100 pM; Probe: mRNA1 FP (650–658) _Middle Region_9 nt_Cy5 at 50 nM; DNA tiles at 6 nM.

**Extended Data Fig. 5.**
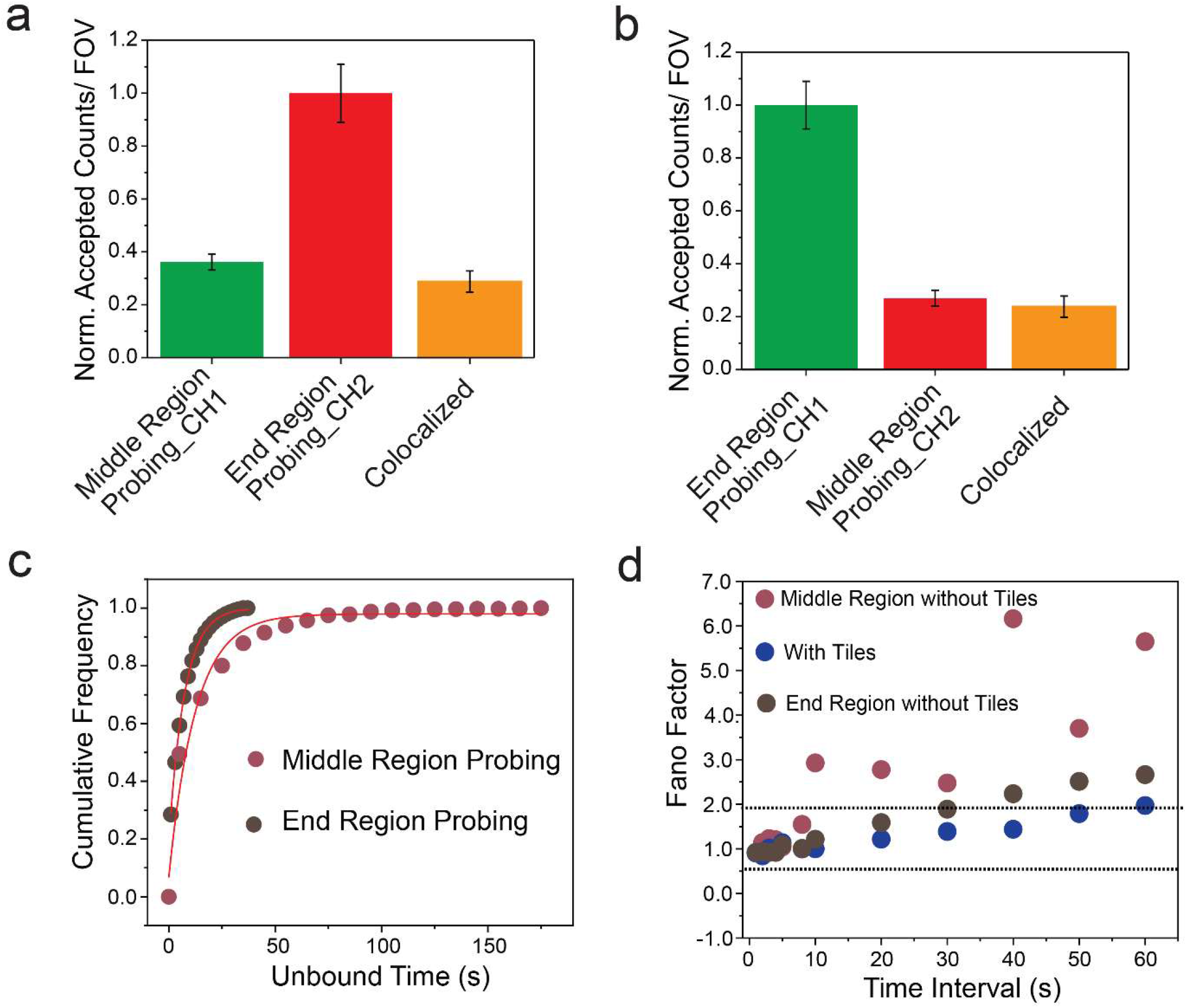
Multi-region single-molecule analysis characterizes regional accessibility and colocalization of encapsulated mRNA1. **a,** Normalized accepted counts per field of view (FOV) for simultaneous probing of encapsulated mRNA1 in the middle region (CH1, Cy3 channel), end region (CH2, Cy5 channel), and colocalized events, without tiles. **b**, Normalized accepted counts per FOV for simultaneous probing where CH1 (Cy3 channel) targets the end region and CH2 (Cy5 channel) targets the middle region, and colocalized counts, without tiles. **c**, Cumulative unbound time histograms comparing middle region and end region probing, with τ_off,med_ values obtained from single-exponential fitting of cumulative unbound time distributions from accepted single-molecule traces. **d**, Fano factor analysis calculated across various time intervals for mRNA1 interactions with specific probe: without tiles and middle region probing (burgundy circles; non-Poissonian), with tiles (blue circles; Poissonian), and without tiles and end region probing (brown circles; intermediate behaviour). Dashed lines show the 95% confidence interval for Poisson statistics. Temperature: 25°C; Acquisition time: 5 min/FOV; Target: LNP1001 at 100 pM; Probe: mRNA1 FP (650–658) _Middle Region_9 nt_Cy5 & Cy3 at 50 nM; mRNA1 FP (2033-2040) _End Region_9 nt_Cy5 & Cy3 at 50 nM; DNA tiles at 6 nM.

**Extended Data Fig. 6.**
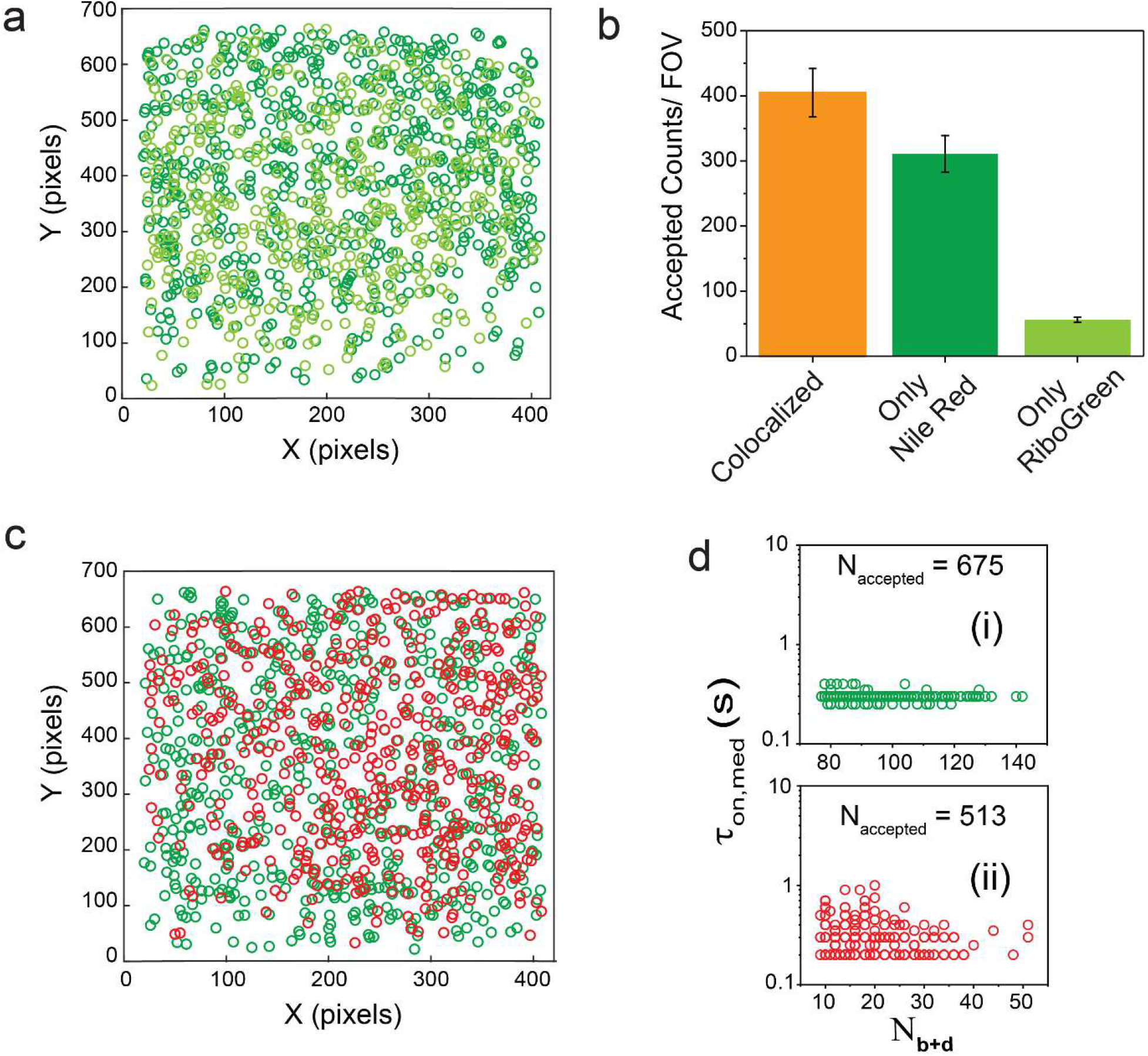
Colocalization-based enumeration of mRNA copies and encapsulation within LNPs using synergized single-molecule assays. **a,** Colocalization analysis of Nile Red (deep green) and RiboGreen (light green) datasets for encapsulated mRNA1 detection inside LNP1001, with both movies acquired sequentially on the Cy3 channel and spot registration maintained. **b**, Accepted counts per field of view (FOV) for encapsulated mRNA1 probed by Nile Red (deep green), RiboGreen (light green), and colocalized events (orange), without tiles. **c**, Colocalization analysis of Nile Red and mRNA1-specific probe datasets for encapsulated mRNA1 inside LNP1001, acquired simultaneously in Cy3 (Nile Red, deep green) and Cy5 (mRNA1-specific probe, red) channels, with spot registration maintained. **d,** (i) and (ii) scatterplots of τ_on,med_ vs N_b+d_ showing accepted counts for Nile Red (deep green) and mRNA-specific probe (red). Temperature: 25°C; Acquisition time: 5 min/FOV; Target: LNP1001 at 100 pM; Probe: Nile Red at 1 nM; RiboGreen at 0.5%; mRNA1 FP (650–658) _Middle Region_9 nt_Cy5 at 50 nM; DNA tiles at 6 nM.

